# *Pseudoarthrobacter* BD237 volatile organic compounds promote *Brachypodium distachyon* growth

**DOI:** 10.64898/2026.10.26.762054

**Authors:** Carl Otto Pille, I Nyoman Sumerta, Hang-Wei Hu, Zahra F. Islam

## Abstract

Microbial volatile organic compounds (mVOCs) present an environmentally sustainable alternative or supplement to traditionally used agrochemicals, such as fertilisers, herbicides and pesticides. Microbial VOCs can promote the growth and stress tolerance of a variety of dicot plants, largely through modulation of phytohormone pathways. However, their effect in monocot plants, particularly grasses, remains largely unexplored. *Pseudoarthrobacter* BD237’s mVOCs were assayed for their growth promoting effect in the model cereal, *Brachypodium distachyon*. BD237 mVOCs were found to significantly increase the dry shoot and root weight of *B. distachyon* by 25% and 42%, respectively. Gas chromatography analysis showed that BD237 produced a variety of alcohols and ketones. Exposure to pure standards of phenylethyl alcohol and 2-nonanone were found to significantly promote *B. distachyon* growth. Transcriptome analysis of *B.distachyon* roots exposed to BD237 mVOC revealed significant upregulation of genes related to photosynthesis and response to abiotic stress. Transcriptome analysis of *BD237* plate colonies exposed to *B. distachyon* VOCs revealed no significant differential transcription, indicating that BD237 VOC production may be constitutive. These results suggest that BD237 mVOCs may promote *B. distachyon* growth by increasing photosynthetic rate and may increase tolerance to abiotic stressors such as salinity and drought. Collectively, this work expands our understanding of microbe – monocot VOC mediated interactions and identifies specific VOCs as potential grass biostimulants.

## Introduction

Agrochemicals such as synthetic fertilisers, herbicides and pesticides can have deleterious effects on environmental and societal health (Elumalai et al., 2025). Thus, sustainable alternatives such as plant growth promoting rhizobacteria (PGPR) and their bioactive compounds are receiving increased interest (Singh, 2025). PGPR inhabit the space on, around or inside plants roots and exhibit multiple, varied plant growth promoting mechanisms. Some improve plant nutritional status by fixing nitrogen, solubilising phosphate or increasing the bioavailability of metals and other micronutrients (Singh et al., 2025). Others produce growth stimulating phytohormones or immune boosting compounds which improve a plant’s tolerance to biotic and abiotic stressors, such as insect herbivory or drought (Singh et al., 2025). However, a large challenge in the application of PGPR is their tendency to be outcompeted by indigenous soil microbiomes and subsequent inability to persist over time (Li et al., 2022). Thus, research has shifted to investigate whether beneficial PGPR produced compounds can be used directly, without the need to rely on the survival of introduced PGPR.

Compared to other plant growth promoting mechanisms, the microbial production of bioactive volatile organic compounds (VOCs) remains relatively unexplored. VOCs are compounds with low boiling points, low vapour pressure and low molecular weights (< 300 Da) (Fincheira and Quiroz, 2018). VOCs are chemically diverse, including alcohols, esters, ketones, pyrazines, aromatics and others, arising from both primary and secondary metabolism (Ryu et al., 2020). Bacterial VOC producers include members of the *Bacillus, Pseudomonas, Pseudoarhtrobacter, Burkholderia, Priestia, Serratia* and *Stenotrophomonas* genera, whilst fungal genera include *Trichoderma* and *Cladosporium* (Ryu et al., 2020, Razo-Belmán et al., 2023). Growth modulating microbial VOCs produced by different bacterial and fungal genera have been shown to significantly increase plant biomass through modulation of photosynthetic rates or phytohormone pathways (Gutiérrez-Luna et al., 2010, Zhang et al., 2008b, Fincheira and Quiroz, 2018). However, a plethora of bioactive mVOCs likely remain unknown, as additional compounds are continuously being discovered in an ever-increasing variety of bacterial strains (Camarena-Pozos et al., 2019) and microbial growth media composition has been shown to have a significant impact on microbe volatilomes (Blom et al., 2011a, Rath et al., 2018).

To date, most of the studies investigating mVOC effects on plant growth have used dicot plants, such as the model plants *Arabidopsis thaliana* and *Nicotiana tabacum*. VOC-mediated growth promotion in dicots has been shown to lead to greater root branching, root length, leaf number, leaf area and higher shoot and root biomass, with gene expression studies in dicots highlighting that mVOC mediated growth primarily linked to phytohormone network modulation, although increased iron uptake and photosynthetic rates have also been observed (Bhattacharyya et al., 2015, Hao et al., 2016, Jiang et al., 2019, Wang et al., 2021, Gao et al., 2022, He et al., 2022, He et al., 2023, He et al., 2024). A recent study also found that a volatile mediated cross talk between *Arabidopsis* and a root bacteria resulted in significant transcriptional changes in the bacterium (Kulkarni, Mazumder et al. 2024).

As monocot plants provide half of the world’s caloric intake (Poutanen et al., 2022), investigation into mVOCs as sustainable alternatives to agrochemicals in monocot cropping systems is highly warranted. Compared to dicots, studies on mVOC mediated growth modulation in monocots are relatively scarce, with no reports on monocot transcriptome responses to mVOCs and few reports on which pure mVOCs stimulate monocot plant growth (Shi, Niu et al. 2017, Paul, Mahmud et al. 2022, Mahadik and Kumudini 2025). Until now, no studies have investigated a monocot plants VOCs effects on a beneficial bacterium which produces plant stimulating mVOCs. Consequently, this study aimed to determine the bioactive effect of mVOCs produced by a root associated bacterial strain on the growth of the model monocot *Brachypodium distachyon,* before identifying bioactive mVOCs and elucidating the dual-transcriptomic response of *B. distachyon* roots to bioactive mVOCs.

## Materials and Methods

### Bacterial strains and culture conditions

*Pseudoarthrobacter* BD237 was previously isolated from sonicated *B. distachyon* roots and identified by 16S Sanger sequencing, as described previously (Pille et al., 2024). BD237 was stored at -80 ℃ in 25% v/v glycerol, and routinely revived and grown on LB liquid and solid medium (10 g/L tryptone, 5 g/L yeast extract, ±10g/L agar).

### In vitro co culturing system

An *in-vitro* co-culture system was adapted from a previous study (Delaplace et al., 2015). A 12 cm square petri dish contained 50 mL of a ¼ Hoagland’s plant agar medium containing macronutrients as follows: 5 mM Ca(NO_3_)_2_, 5 mM KNO_3_, 2 mM MgSO_4_, 41.5 µM FeEDTA(Na), 1 mM KCl, 1 mM KH_2_PO_4_ and micronutrients as: 92 µM H_3_BO_3_, 20 µM MnCl_2_, 1.6 µM ZnCl_2_, 0.5 µM CuCl_2_ and 0.24 µM Na_2_MoO_4_ with 0.8% (w/v) agar (Hoagland and Arnon, 1950).

Hoagland’s agar medium was buffered to pH 6.5 with 1 mM MES-hydrate. In the middle of the square petri dish, a smaller, round petri dish (3.5 cm diameter) containing 3 mL of either LB agar or ½ MS agar, (both with 1.5% (w/v) agar) was placed while the plant growth agar was still molten. In this *in vitro* system, plants and bacteria were physically separated (Fig. 1).

**Figure 1:**
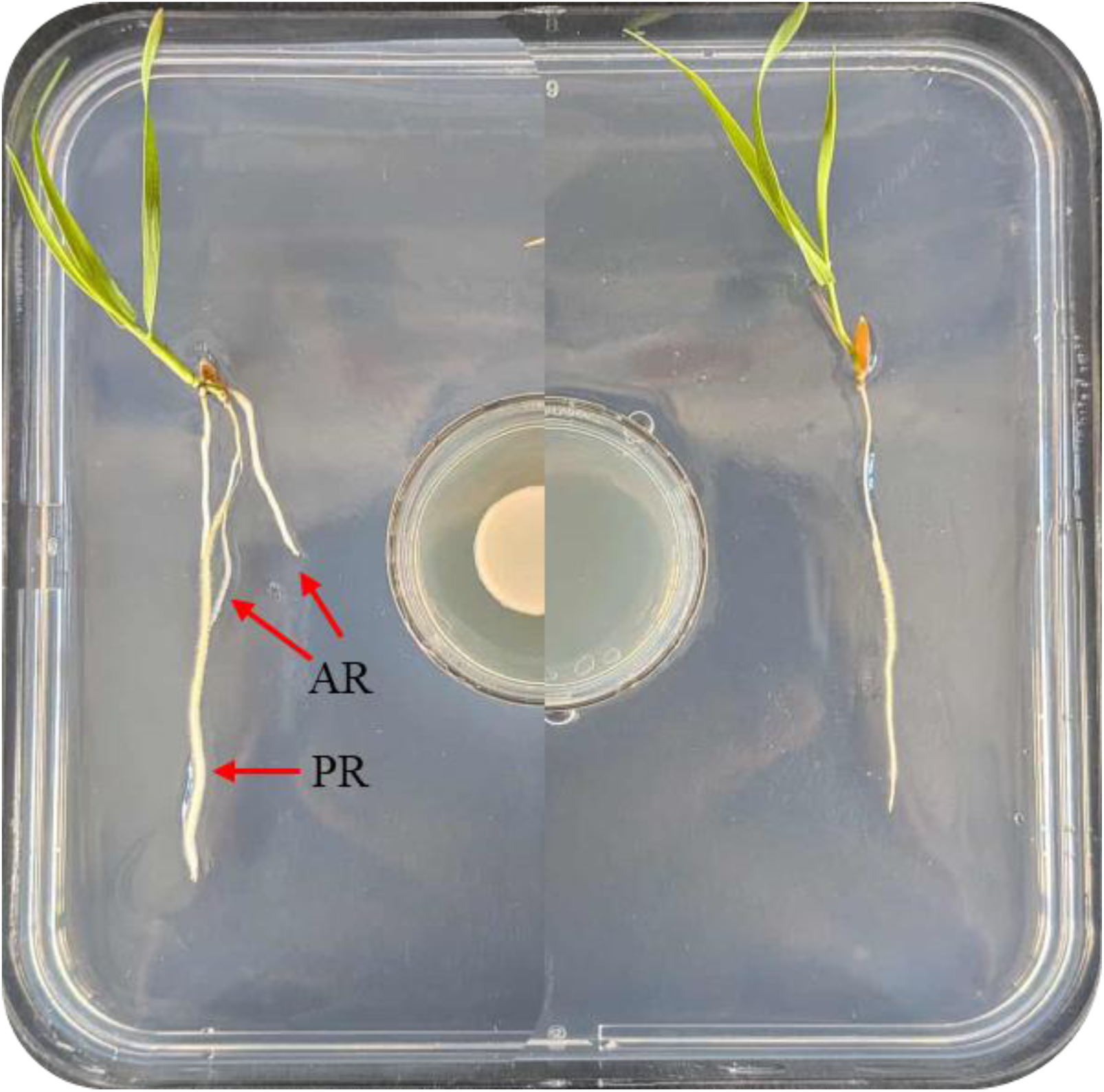
Co-culture system used to investigate potential growth modulating effect of bacterial VOCs on *B. distachyon* growth. Representative plantlets were co-cultivated for 10 days with (right) or without (left) exposure to *Pseudoarthrobacter* BD237 volatiles. Red arrows indicate primary root (PR) and adventitious roots (AR).

Dehusked and surface sterilised *B. distachyon* seeds were germinated on 0.9% water germination agar at room temperature for 48 h. Seedlings with a similar radicle length (∼ 1.5 cm) were selected for experimental use. *Pseudoarthrobacter*BD237 were plated onto LB agar plates and incubated for 48 h at 28 ℃. Single colonies were picked from LB plates and used to inoculate liquid LB. Liquid LB cultures were grown for 24 h at 28℃ with shaking at 250 rpm. Liquid cultures were then adjusted to an OD_600_ equivalent to approximately 10^8^ CFU/mL (OD_600_ ∼ 0.5). Two seedlings were placed towards the top of each plate with their embryos facing the plate’s lid. Subsequently, 20 µL of each bacterial culture were spotted onto the middle of the bacterial growth compartment and dried in a laminar flow hood. Square plates were then sealed with parafilm and placed at a 65° angle to ensure roots grew on the surface of the growth agar (Delaplace, Delory et al. 2015). Plates were then incubated in a Conviron growth cabinet under a 18/6h light cycle at 24℃ and 18℃, respectively. Plants and bacteria were co-cultured for 10 days before harvesting. Six plates with a total of twelve seedlings were included for each treatment and control condition. Significant differences between control and treatments were determined using one-way ANOVA and post-hoc Tukey tests.

### *B. distachyon* growth modulation by pure VOCs

Plants were prepared as described previously for assessment of growth modulation by pure VOCs. Instead of filling the small, middle compartment with LB agar, a 1.5 cm filter paper was soaked with 20 µL DMSO containing 2, 20, 200 or 2000 ng of selected VOCs. Control condition consisted of DMSO only. Plants and bacteria were co-cultured for 10 days before harvesting. Four plates with a total of 8 seedlings were included for each treatment and control condition. Significant differences between treatments were determined via one-way ANOVA and post-hoc Tukey tests.

### Root system architecture analysis

After 10 days of co-culture, each plantlet had its roots untangled using a dissecting needle and tweezers before placing onto 0.9% 12 cm square agar plates and scanned using an Epson Perfection V800 scanner at 600 dpi. Image artefacts in scans were manually corrected using Microsoft Paint before analysis using the open-source software RhizoVision Explorer (Fig. S4.1) (Seethepalli et al., 2021).

### Growth modulation by mVOCs in an *in vitro* soil system

Magenta boxes were modified to investigate the effect of mVOCs on *B. distachyon* plants grown in soil. Boxes used to hold soil and *B. distachyon* had nine evenly spaced 6mm holes drilled into the bottom. A 6 x 6 cm square Whatman #1 filter paper was placed at the bottom of the box to prevent soil escaping. *B. distachyon* seeds were prepared and germinated as described previously. Two-day old plantlets were then sown into Magenta boxes containing Plugger™ seed raising mix. Seedlings were grown to the two-leaf stage (5 days) before introducing contact to mVOCs.

*Pseudoarthrobacter* BD237 was revived and cultured as described above and a 20 µL droplet spotted on to the centre of a 3.5 cm petri dishes containing 3 mL LB agar. Bacterial droplets were dried under laminar flow before sealing with Parafilm and placing inside Conviron growth cabinets under the same conditions described previously. Isolates were grown for 48 h before 9 holes were melted into the lids of their plates using a flamed dissection needle inside a laminar flow hood. Bacterial plates were then immediately placed at the bottom of unmodified Magenta boxes. Modified Magenta boxes containing soil and *B. distachyon* plantlets were then placed inside Magenta boxes containing bacterial plates before sealing their junction with Parafilm (Fig.3a). Plants and bacteria were allowed to grow together for 10 d before harvesting and recording plant growth parameters.

### mVOC identification via SPME-GC-MS

*Pseudoarthrobacter* BD237 was plated onto LB agar plates and grown for 48 h at 28 ℃. Single colonies from these plates were then picked and used to inoculate liquid LB. Liquid cultures were grown overnight and adjusted to an OD_600_ of 0.5 (CFU ∼ 1x10^7^). A 20 µL droplet of each inoculant was then spotted onto 9 mL of LB agar in a sterile 20 mL glass gas chromatography vial and allowed to dry before closing the vial with its surface sterilized screw top lid. GC vials containing bacteria (and controls containing only LB agar) were then placed in a Conviron growth cabinet under identical conditions used during *in vitro* growth promotion assays and incubated for 4 d. The headspace of incubated GC vials were then analysed via solid phase microextraction and gas chromatography coupled to a mass spectrometer (SPME-GC-MS/MS). In short, a 10 mM (Supelco®, Sigma Aldrich) coated with 65 µm poly(dimethylsiloxane) and divinylbenzene (PDMS/DVB) was inserted into sample vials and volatiles allowed to adsorb for 30 min at 50 ℃. Fibres were then desorbed into the injection port of an Agilent 8890A GC system with an Agilent DB5 column (30 m × 0.25 × 0.25 µm film thickness) coupled to an Agilent 59778B MS (Agilent Technologies) and equipped by an Agilent PAL RSI 85 autosampler. Samples were run according to previously reported parameters (Farag et al., 2006) and peaks were identified by comparison to the National Institute of Standards and Technology database (Kramida, 2024).

### *B. distachyon* root transcriptome analysis in response to BD237 VOCs

*B. distachyon* plantlets and *Pseudoarthrobacter*BD237 were co-cultured in an identical manner to the *in vitro* co culturing system described previously. After 10 days, basic plant parameters were recorded and roots were removed from one plantlet per plate, with five plates each for control and treatment conditions. Roots were then immediately frozen in liquid nitrogen (LN_2_) before grinding into a powder in an LN_2_ cooled, autoclave sterilised mortar and pestle. A QIAGEN RNeasy Mini Kit was then used to extract RNA from root powders, as per manufacturer’s instructions, except that a lower biomass was used (∼ 50 mg) than recommended, as a higher biomass led to over saturation of the kit’s columns. RNA samples were then digested with DNase to remove any contaminating DNA, before sending samples to AGRF in Melbourne, Victoria for Illumina RNA sequencing.

The following pre-processing steps were conducted in the Galaxy Australia web platform (2024). Reads from triplicate lanes per sample were concatenated. Raw reads qualities were assessed with Falco, before trimming of low-quality bases and adapter sequences using Trim Galore! (de Sena Brandine and Smith, 2021). The cleaned data were then aligned to the EnsemblPlants *Brachypodium distachyon* v3.0 reference genome (https://plants.ensembl.org/Brachypodiumdistachyon/Info/Index) using the HISAT2 software v2.2.1 (Kim et al., 2015a). The featureCounts software v2.1.1 was used to assign mapped reads to exons using the EnsemblPlants v3.0.61 annotation file (https://ftp.ensemblgenomes.ebi.ac.uk/pub/plants/release62/gff3/brachypodiumdistachyon/) (Liao et al., 2014).

The following steps were conducted in R studio (R Core Team, 2013). DESeq2 v2.11.40.8 was used to find genes differentially expressed between control and treatment samples (Love et al., 2014). P-values were adjusted using the Benjamini-Hochberg procedure. A list of all expressed genes were ranked by Log2 Fold Change and used to conduct Gene Ontology (GO) pathway enrichment analysis using the GSEA function in the clusterProfiler package v4.16.0 (Yu et al., 2012).

### *Pseudoarthrobacter* BD237 genome assembly, annotation and phylogenetic analysis

A single BD237 colony grown overnight on an LB agar plate at 28℃ was picked and used to inoculate an overnight liquid LB culture grown at 28℃ and 250 rpm. Aliquots of the liquid culture were then centrifuged for 10 mins at 5,000 × g to pellet bacterial cells. Genomic DNA was extracted from bacterial pellets using the Promega Wizard Genomic DNA Purification Kit, per manufacturer instructions. Genomic DNA was quantified via Nanodrop and then sent to AGRF in Melbourne, Victoria for quality control and subsequent Illumina short-read DNA sequencing in paired end mode.

The following steps were conducted in the Galaxy Australia web platform. Raw read quality was assessed with Falco (Version 1.2.4), before trimming low quality (Phred score < 30) bases and adapter sequences using Trim Galore! (Version 0.6.10) (de Sena Brandine and Smith, 2021). *De novo* assembly of the BD237 genome was conducted using the Shovill pipeline (Version 1.4.2; https://github.com/tseemann/shovill). Contigs smaller than 1000 base pairs were filtered out. Assembly completeness was determined using QUAST (version 5.3.0), BUSCO (version 5.8.0) and CheckM2 (version 1.1.0) (Gurevich, Saveliev et al. 2013, Simão, Waterhouse et al. 2015). The remaining assembled contigs were annotated using the Bakta pipeline (Version 1.9.4) (Schwengers, Jelonek et al. 2021). Annotation of CDS regions with Gene Ontology (GO) terms was then performed using EggNOG Mapper (version 2.1.13) (Huerta-Cepas, Forslund et al. 2017).

Relationships to other microorganisms were estimated based on average nucleotide identity (ANI) and DNA-DNA hybridisation (dDDH) rates between the BD237 assembly and the closest relatives using the FastANI tool and the Type Strain Genome Server online portal using default parameters, respectively (https://tygs.dsmz.de/) (Freese, Meier-Kolthoff et al. 2026). ANI was calculated using default parameters against 68 reference genomes from the genera *Arthrobacter*, *Pseudoarthrobacter* and *Paenoarthrobacter*. Of these, 65 were type strain assemblies and 2 were no type strains for *P. niigatensis*, for which no type strain exists. Finally, a phylogenetic tree was inferred from GTDB-Tk v2.6.1 generated concatenated alignments of 120 marker genes present in the previously mentioned reference genomes. The IQ-TREE tool was used to generate a maximum likelihood phylogenetic tree, with branch support estimated from SH-aLRT (Shimodaira-Hasegawa-like approximate likelihood ratio) tests and 1000 ultrafast bootstrap replicates. Nodes were considered as well supported when SH-aLRT and ultrafast bootstrap values were ≥ 80% and ≥ 95%, respectively (met by 48 of 66 nodes). The tree was rooted on the genus *Paenoarthrobacter*.

### BD237 transcriptome analysis in response to *B. distachyon* VOCs

*B. distachyon* plantlets and *Pseudoarthrobacter*BD237 were co-cultured in an identical manner to the *in vitro* co culturing system described previously. After 10 days, BD237 colony edges were collected and RNA was extracted using the QIAGEN All Prep DNA/RNA/Protein kit, as per the manufacturer’s instructions. RNA samples were then digested with DNase to remove any contaminating DNA, before sending samples to AGRF in Melbourne, Victoria for ribosomal RNA depletion and Illumina short read RNA sequencing.

The following steps were conducted on the Galaxy Australia platform. Raw reads qualities were assessed with Falco, before trimming of low-quality bases and adapter sequences using Trim Galore! (de Sena Brandine and Smith, 2021). The cleaned data were then aligned to the BD237 genome assembly using HISAT2. Aligned reads were then assigned to genes using the BD237 genome previously annotated using the Bakta tool. The following steps were conducted in R studio (R Core Team, 2013). DESeq2 v2.11.40.8 was used to find genes differentially expressed between control and treatment samples (Love et al., 2014).

## Results

### Phylogenetic placement of BD237

The draft genome of BD237 comprised 51 contigs with a total length of 4,668,822 bp, a GC content of 66.6% and an N50 of 184,180 bp. CheckM2 estimated the assembly to be 99.99% complete with 0.30% contamination and BUSCO v5.8.0 recovered 99.5% of the 1050 single-copy orthologues of the *Arthrobacter* lineage dataset, strongly indicating genome completeness. Annotation with Bakta v1.9.4 predicted 4407 genes, comprising 4346 protein-coding sequences, 52 tRNA genes, one tmRNA gene, three ncRNA genes and four rRNA genes.

Whole genome and phylogenetic analyses indicate that strain BD237 is a distinct species, most closely related to *Pseudoarthrobacter niigatensis* (Fig. S1). Overall genome relatedness indices (ANI & dDDH) place BD237 below both species boundaries against every genome examined: ANI to the *P. niigatensis* genomes was 92.73–92.76%, with no reference among 68 reaching the 95% threshold, while the Digital DNA-DNA results showed 48.5% dDDH with *P. niigatensis*, well below the 70% dDDH species threshold (Fig. S1).

### *Pseudoarthrobacter* BD237 mVOC mediated growth modulation in *B. distachyon*

VOCs produced by *Pseudoarthrobacter* sp. BD237 growing on LB media significantly increased *Brachypodium distachyon* shoot and root dry weight by 25% and 42%, respectively (Fig. 2a, c). VOCs produced on LB media also significantly increased *B. distachyon* root length and number of adventitious roots (Fig. 2b, d). As the negative control strain *E. coli* DH5α did not promote plant growth in additional assays, the effect of microbial respiration (production of CO_2_) was deemed to not significantly affect plant growth in the co-culture system (Fig. S2). When cultured on ½ MS media, BD237 VOCs led to a non-significant reduction in shoot and root biomass in *B. distachyon* plantlets (Fig. 2a, c). After exposure to BD237 mVOCs, soil grown *B. distachyon* plantlets’ shoot dry weight increased significantly by 19% (Fig. 3b).

**Figure 2:**
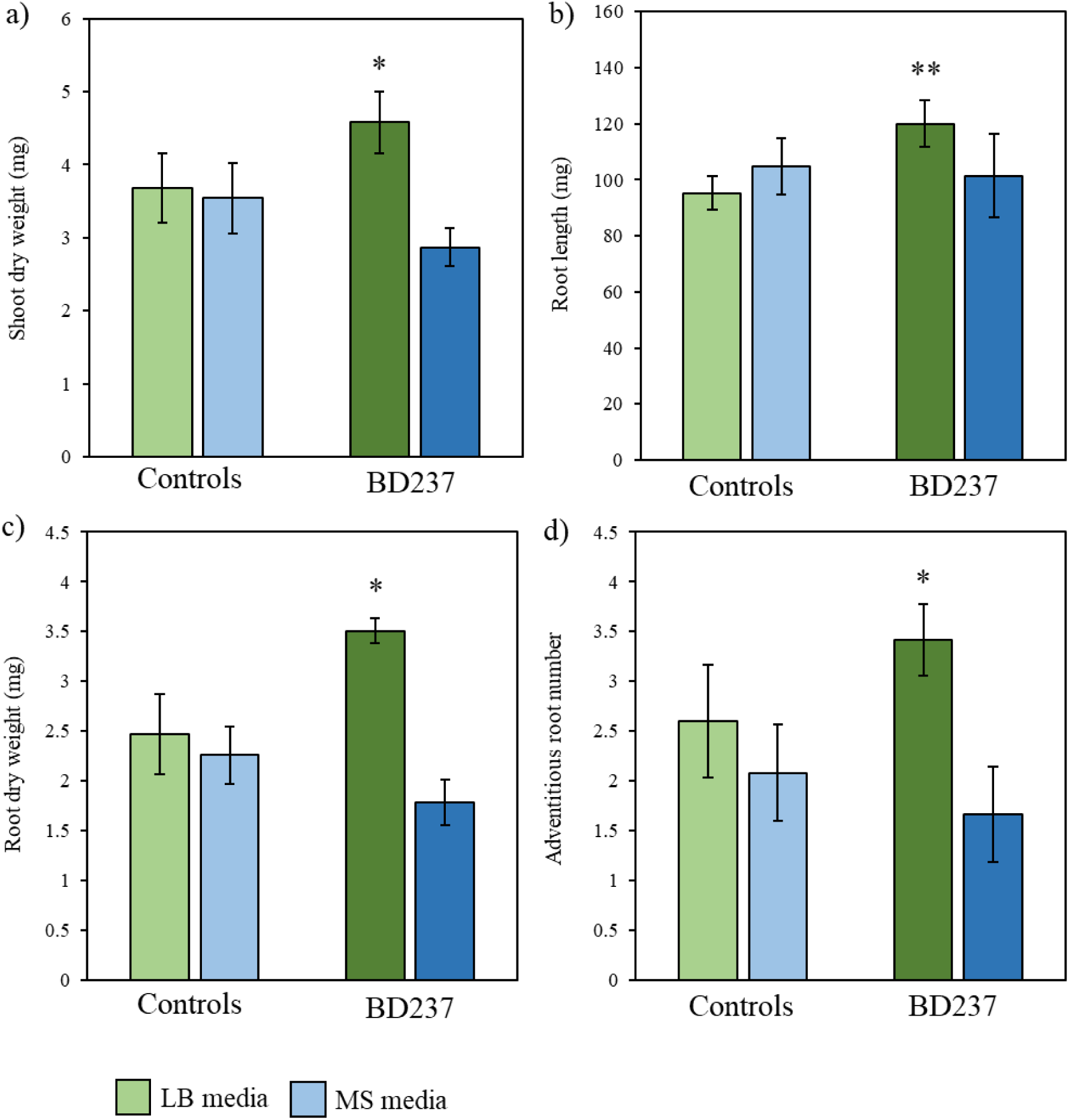
Mean dry shoot (a) and root (b) weights of *B. distachyon* plantlets exposed to BD237 VOCs for 10 days with associated 95% confidence intervals (* = *p* < 0.05, ** = *p* < 0.01, *** = *p* < 0.00, 1 n = 12). Green bars indicate plants exposed to volatiles of bacteria grown with LB, blue bars indicate plants exposed to volatiles of bacteria grown with ½ MS. Mean adventitious root number (c) and total root length (d) of *B. distachyon* plants exposed to BD150 and BD237 volatiles for 10 days with associated 95% confidence intervals (* = *p* < 0.05, ** = *p* < 0.01, n = 12)

**Figure 3:**
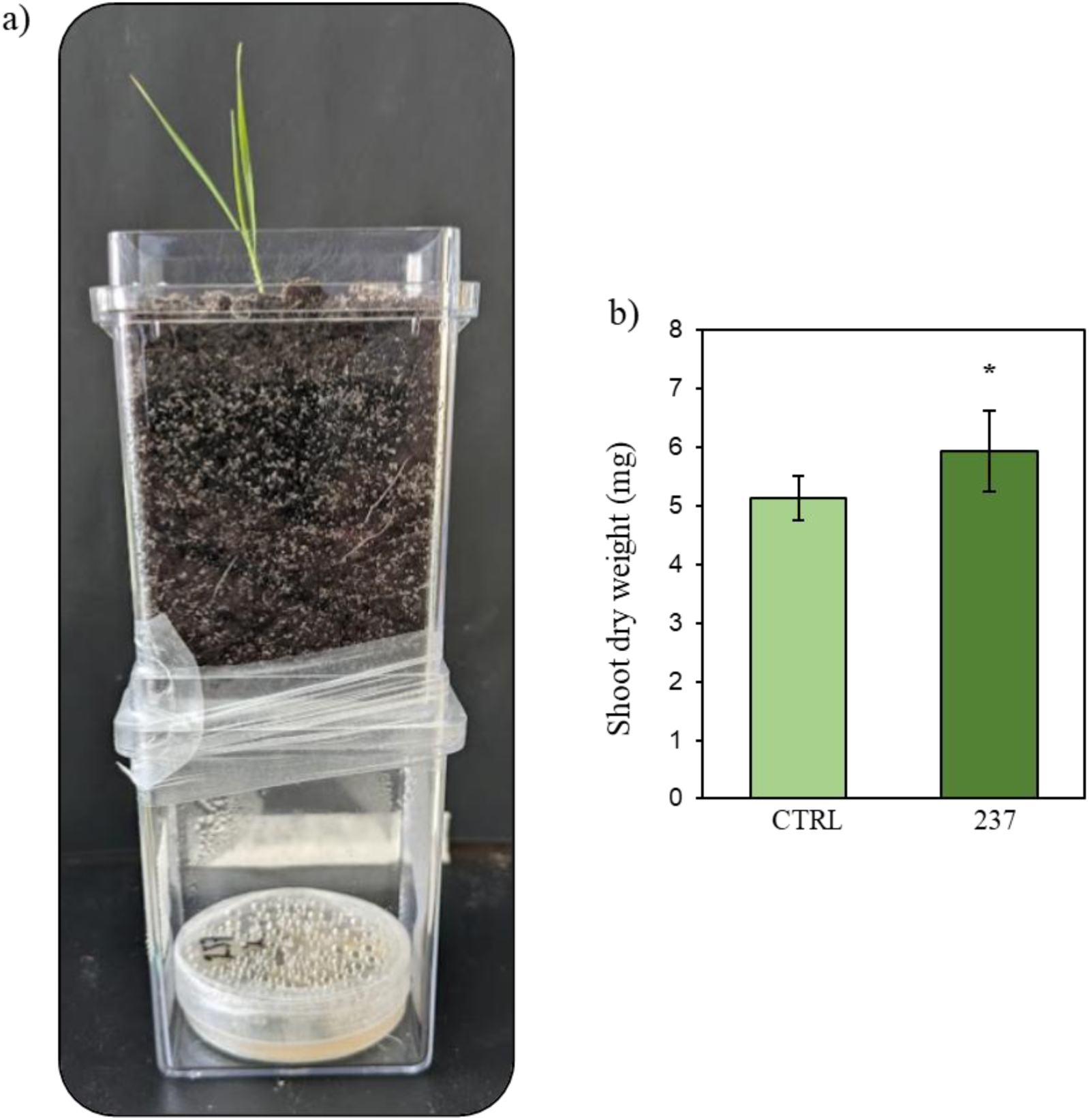
(a) *In vivo* co-culture of *B. distachyon* in soil (top) and *Pseudoarthrobacter*BD237 culture (bottom). Holes in the bottom of the top compartment and the lid of BD237 plate cultures allow microbial VOCs to reach plant roots. (b) Mean shoot dry weight of *B. distachyon* plantlets grown in soil and exposed to BD237 VOCs for 10 days with associated 95% confidence intervals (* = *p* < 0.05, ** = *p* < 0.01, n = 5).

### VOCs produced by *Pseudoarthrobacter*BD237 on LB agar

SPME-GC-MS was used to identify BD237 mVOCs produced when grown on LB agar, as growth on this media led to the production of plant beneficial mVOCs. SPME-GC-MS results show that *Pseudoarthrobacter* BD237 produces a range of VOCs when grown on LB agar. VOC classes produced by BD237 include ketones, alcohols and pyrazines (Table 1). Subsequently, phenylethyl and benzyl alcohols were the most relatively abundant detected VOCs, while the ketones 2- nonanone and 2-dodecnaone were the least relatively abundant.

**Table 1:** VOCs produced by *Pseudoarthrobacter* BD237 grown on LB agar and detected via SPME-GC-MS

| ID (NIST) | Chemical<br>Class | Retention<br>Time (min) | Peak area |
| --- | --- | --- | --- |
| 1-Butanol | Alcohol | 3.4 | 62,363 |
| 1-Butanol, 3-methyl- | Alcohol | 4.8 | 105,493 |
| 2 – heptanone | Ketone | 7.9 | 12,860 |
| Cyclopentanone, 3,4-bis(methylene)- | Ketone | 9.4 | 32,163 |
| Benzyl alcohol | Alcohol | 10.5 | 36,400 |
| Pyrazine, 2-methyl-5-(1-methylethyl)- | Pyrazine | 10.9 | 17,824 |
| 2 - Nonanone | Ketone | 11.5 | 6,385 |
| Phenylethyl alcohol | Alcohol | 11.9 | 140,289 |
| Phenol, 2,5-bis(1,1-dimethylethyl) | Phenol | 17.4 | 31,674 |
| 2-Dodecanone | Ketone | 18 | 5,104 |

Retention time refers to the amount of time an analyte takes to travel from the GC injection port to the mass detector.

### Effect of pure VOCs on *B. distachyon* growth

When applied in their pure forms, several mVOCs identified via SPME-GS were observed to significantly increase *B. distachyon* shoot and root dry weight, as well as significantly increase total root length (Fig. 4). In particular, it was observed that 2000 ng of 2-nonanone significantly increased *B. distachyon* dry shoot and root biomass by 37% and 55%, respectively (Fig. 4a, b). Two, 20 and 2000 ng of 2-nonane also significantly increased *B. distachyon* total root length by 74%, 77% and 65%, respectively (Fig. 4c).

**Figure 4:**
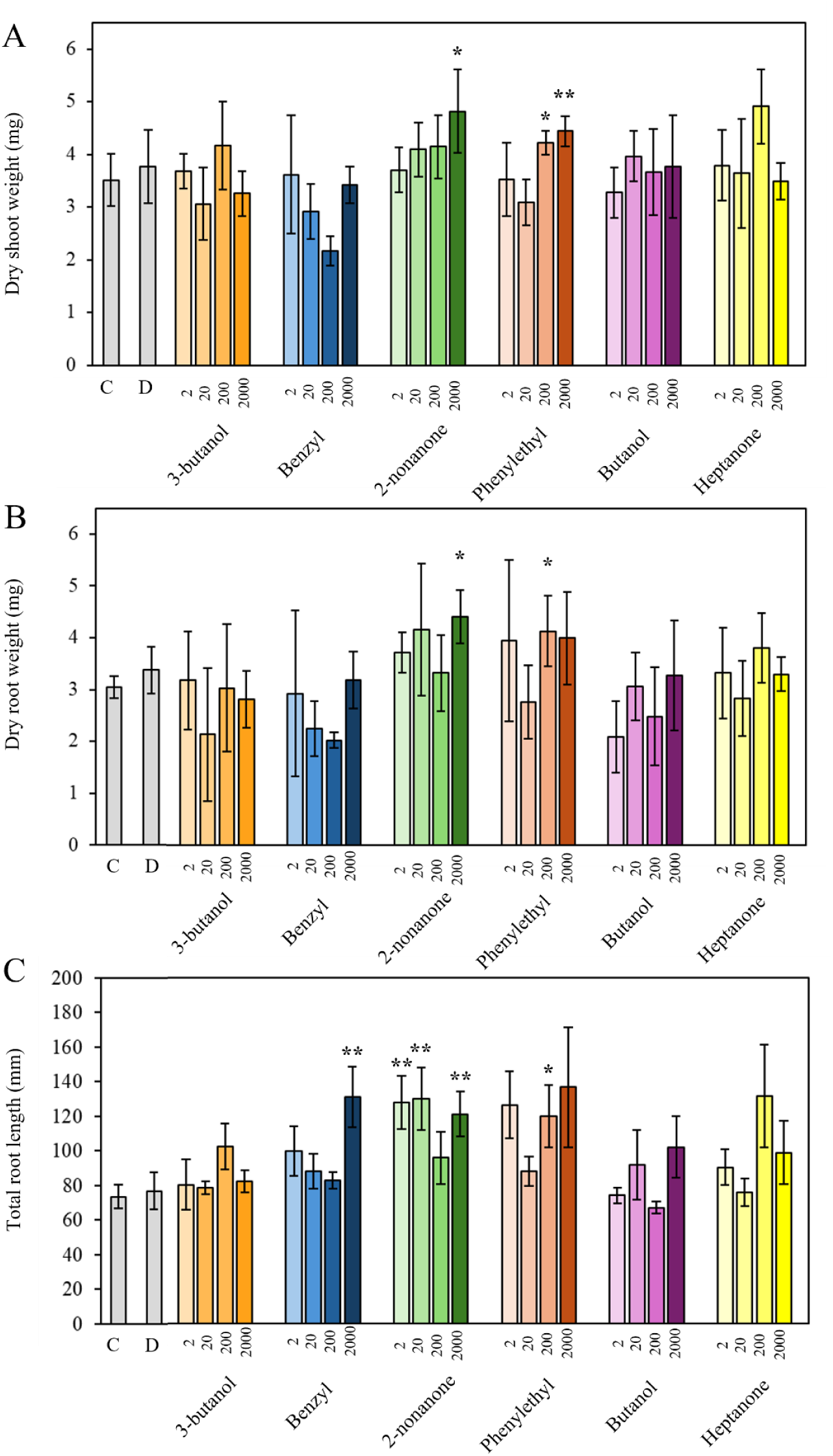
Mean dry shoot (A) and root (B) weights and total root length (C) of *B. distachyon* plantlets exposed to increasing amounts of pure VOCs for 10 days with associated 95% confidence intervals (* = *p* < 0.05, ** = *p* < 0.01, n = 8). Dosages for each VOC range from 2, 20, 200 and 2000 ng (left to right). C = H_2_O negative control; D = DMSO vehicle control.

Additionally, 200 ng of phenylethyl alcohol significantly increased total root length, shoot dry weight and root dry weight by 63%, 20% and 45%, respectively. Similarly, 2000 ng of phenylethyl alcohol significantly increased shoot dry weight by 27% but did not significantly increase root dry weight or total root length (Fig. 4). The vehicle control DMSO, which was used to dilute pure VOC standards, did not modulate *B. distachyon* growth, nor was there an effect of the negative control H_2_O on *B. distachyon* growth.

Retention time refers to the amount of time an analyte takes to travel from the GC injection port to the mass detector.

### *B. distachyon* root transcriptional response to *Pseudoarthrobacter*BD237 volatiles

To determine the transcriptional response to BD237 VOCs, we conducted transcriptome analysis on *B. distachyon* roots. After removal of adapter sequences and low-quality reads, an average of 24.2 million reads remained in each sample. Reads were mapped to the reference genome using HISAT2, with an average mapping rate of 96.8% (Table S1). Mapped reads were then assigned to exonic regions of the *B distachyon* genome via the featureCounts software, with an average of 80.6% of mapped reads assigned (Table S1). Counts were normalised using variance stabilising transformation (VST) via DESeq2. Normalised counts were used to generate a principal component analysis (PCA) plot showing correlation and clustering between samples (Fig. 5). From the PCA plot, it can be observed that mVOC treated samples cluster together and away from control samples, which also cluster together (Fig. 5a). Comparatively, there were no effects of *B. distachyon* VOCs on BD237 transcriptional profiles? (Fig. 5b).

**Figure 5:**
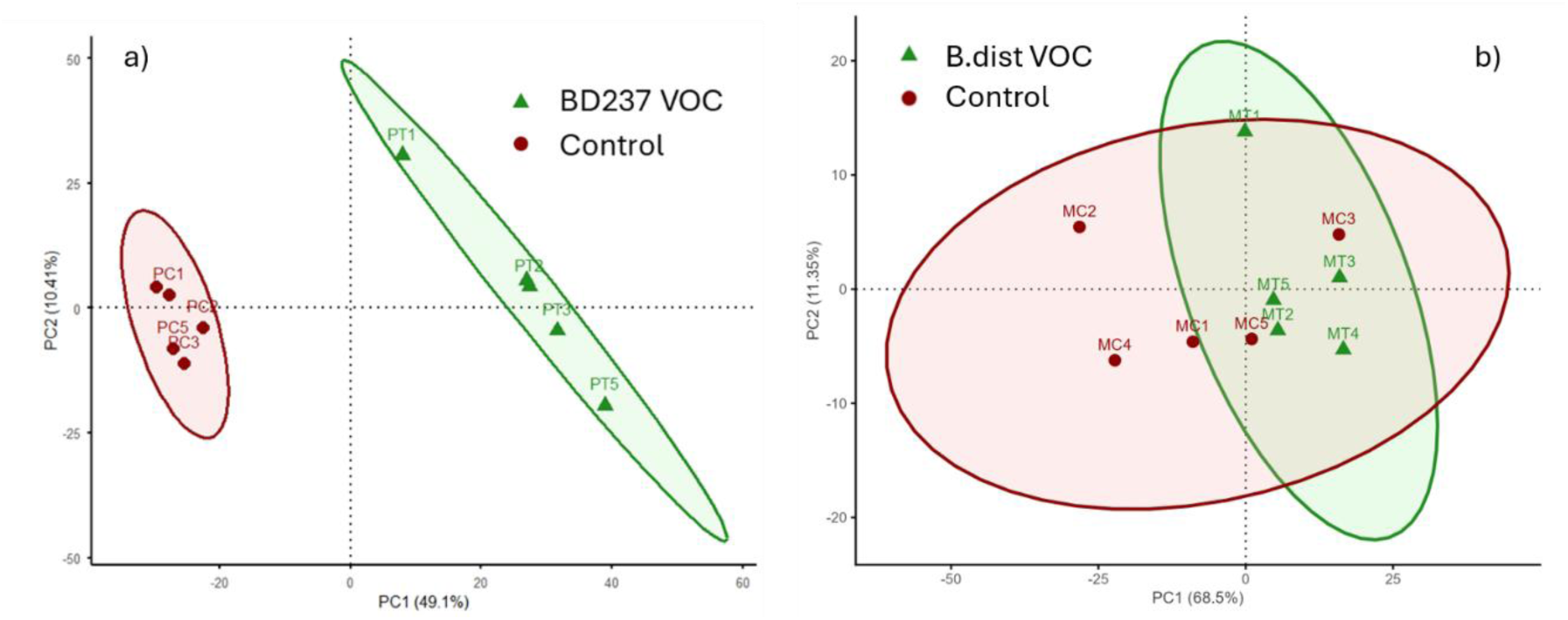
Principal component analysis plots of *B. distachyon* (a) and BD237 (b) RNAseq samples.

Next, DESeq2 was used to determine fold changes in detected genes between control and treatment samples. A total of 29,399 genes were detected and the output of DESeq2 were ranked in descending order according to their log2 fold change. This ranked list was used to determine enrichment of biological pathways in the GO database. Normalised enrichment score plots show where genes related to certain pathways were placed in the ranked list (Fig. S3).

From these analyses, it was observed that exposure to BD237 VOCs upregulated genes related to the GO terms “photosynthesis”, “nutrient reservoir activity”, “response to water deprivation” and “response to abscisic acid”, while downregulated genes related to the terms “RNA modification”, “nucleolus” and “nitrate assimilation” (Fig. 6).

**Figure 6:**
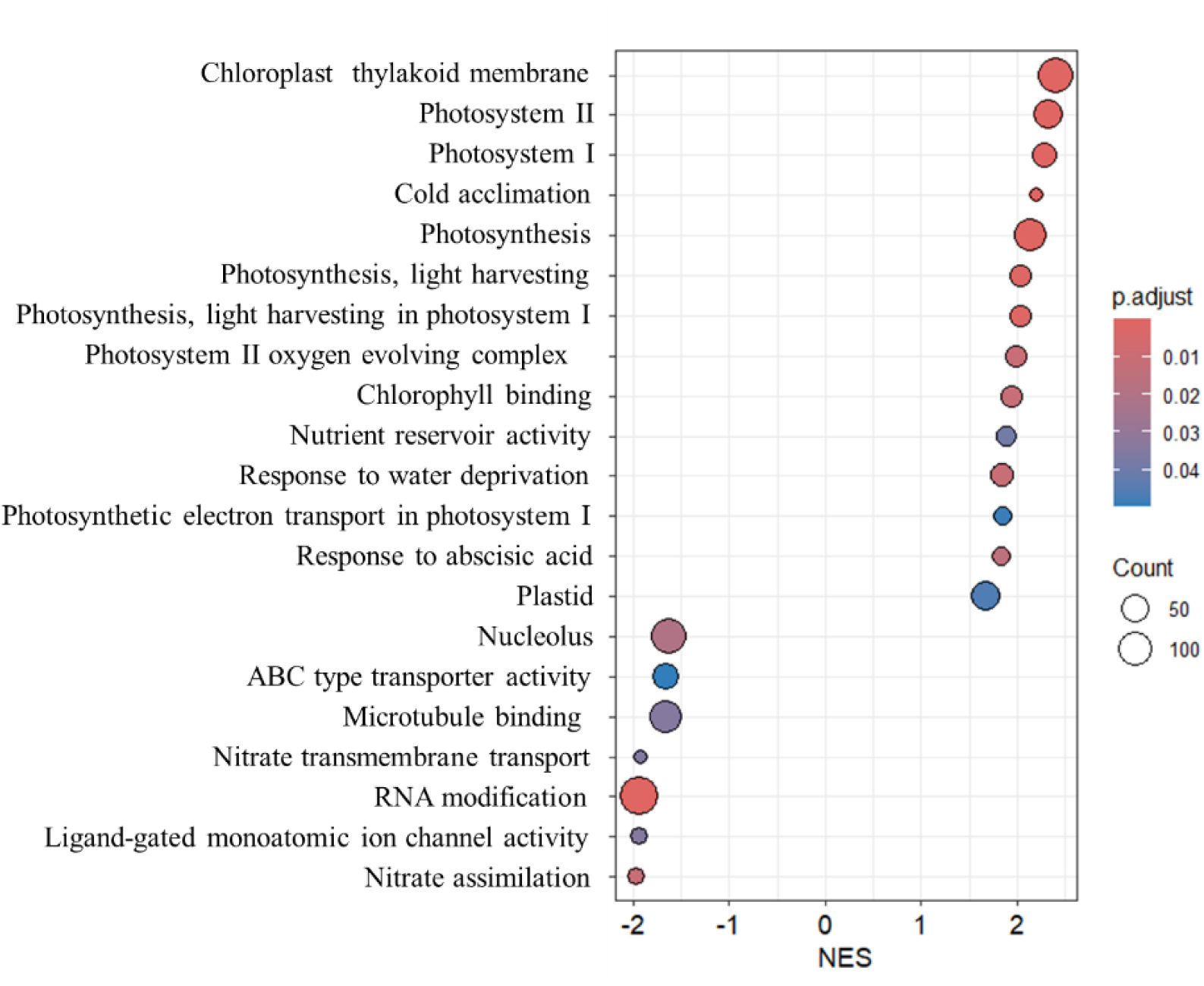
Enriched GO pathways, identified via gene set enrichment analysis using the ClusterProfiler package. The size of the circles corresponds to the number of genes contributing to up or down regulation of each pathway. Colours of circles indicate the significance of pathway modulation.

Although a list of all detected genes ranked by their fold change was used to determine modulated pathways, significantly differentially expressed genes (DEGs) were defined as those with a |log2(Fold Change)| > 1 (2-fold change) and padj < 0.05 to identify genes with large impacts on GO pathway expression. A list of the 20 most significantly up and down regulated genes can be found in Supplementary Tables 2 and 3. Upregulated DEGs in the GO “photosynthesis” pathway included RuBisCo (RBCS) and its subunit (RBCS1), as well as other essential proteins required for photosynthesis (Fig. 7c). Catalase 1 (CAT1), dehydrin 3 (DHN3), Rab21 and MYB2 genes were also significantly upregulated, contributing to enrichment of the “response to abscisic acid” and “response to water deprivation” GO pathways (Fig. 7a, b). In the nitrate assimilation GO pathway, several genes were significantly downregulated, including two copies of the NRT2.1 nitrate transporter, two copies of the NIA1 nitrate reductase, NIR1 nitrite reductase and NRT2.3 nitrate transporter (Fig. 7d).

**Figure 7:**
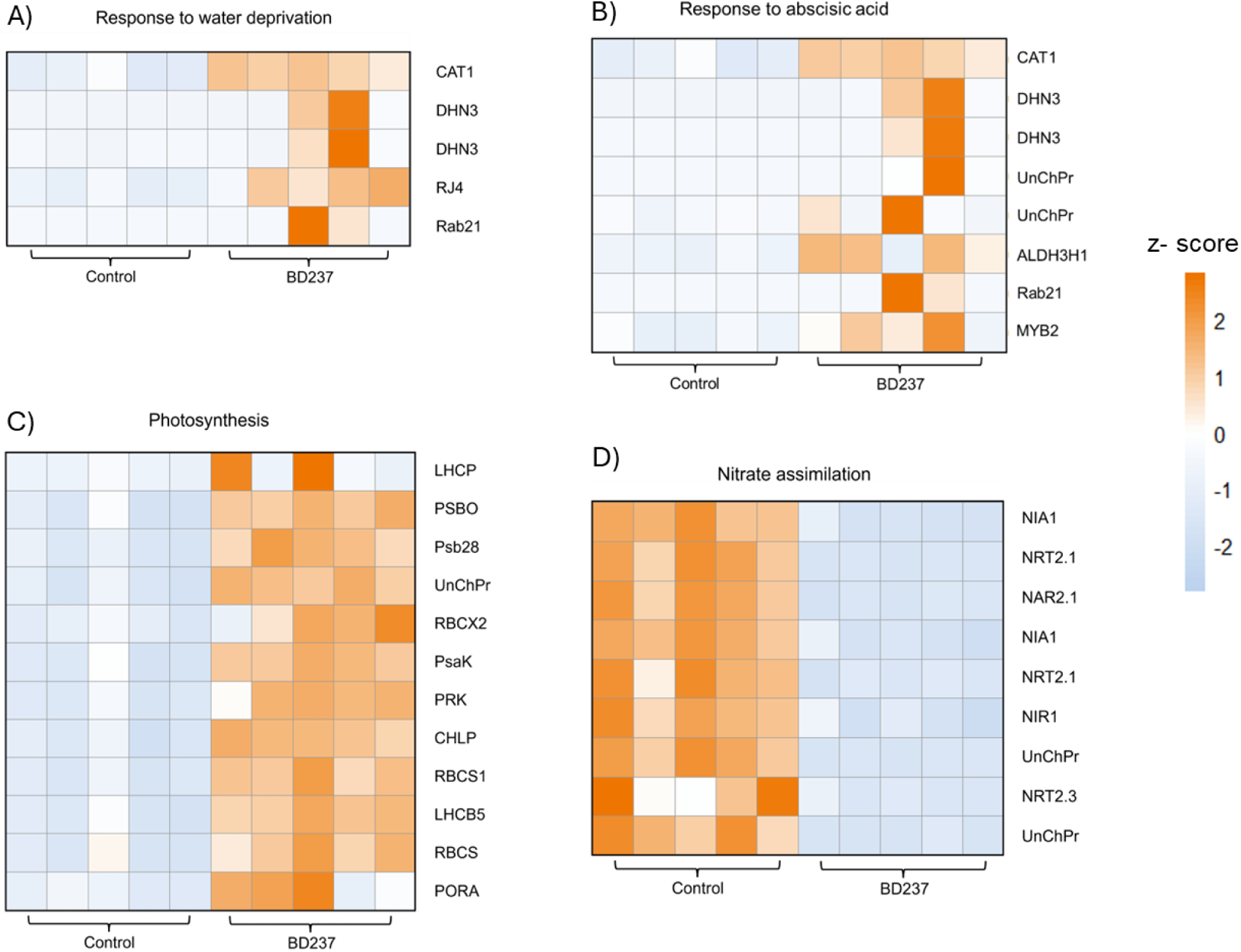
Heatmaps showing expression of *B. distachyon* genes related to up (A, B & C) and down (D) regulated GO terms in response to BD237 mVOCs. Colours of cells are determined by sample z-scores, or number of standard deviations from mean gene expression.

### *Pseudoarthrobacter* BD237 transcriptional response to *B. distachyon* volatiles

To determine the transcriptional response of BD237 to *B. distachyon* volatiles, agar plate colonies were exposed to whole plant volatilomes for 10 days. After removal of adapter sequences and low-quality reads, an average of 18.3 million reads remained in each sample. Reads were mapped to the reference genome using HISAT2, with an average mapping rate of 99.1% (Table S1). Mapped reads were then assigned to exonic regions of the BD237 draft genome via the featureCounts tool, with an average of 75.4% of mapped reads assigned (Table S1). Counts were normalised using variance stabilising transformation (VST) via DESeq2 and used to create a PCA plot to show sample clustering. From this, it was observed that VOC exposed (treatment) and unexposed (control) samples clustered together. Significantly differentially expressed genes were analysed using DESeq2, which showed that no genes met the threshold for significantly differential expression. Overall, exposure of BD237 colonies to *B. distachyon* VOCs did not elicit a detectable transcriptional response.

## Discussion

This study has characterised the effect of *Pseudoarthrobacter* BD237 mVOCs on the growth of the model grass *B. distachyon*. The mVOCs 2-nonanone and phenylethyl alcohol were shown to significantly increase *B. distachyon* biomass and root length. Subsequently, an RNAseq experiment showed that BD237 mVOCs stimulate the upregulation of photosynthesis and stress response related genes, suggesting that BD237 mVOCs promote *B. distachyon* growth by altering photosynthetic rates, while potentially also promoting tolerance against abiotic stressors. However, this effect was not bi-directional; transcriptomic profiling of BD237 exposed to *B. distachyon* VOCs showed no differentially expressed genes.

### PseudoarthrobacterBD237 mVOCs promote B. distachyon growth

Exposure to *Pseudoarthrobacter*BD237 mVOCs led to a significant increase in *Brachypodium distachyon* shoot and root dry weight by 25% and 42%, respectively (Fig. 2a, c). BD237 also increased total root length and number of adventitious roots (Fig. 2b, d). To date, a single other *Pseudoarthrobacter* strain’s VOC has been reported to modulate plant growth via action of the mVOC dimethylhexadecylamine (Velázquez-Becerra et al., 2011, del Carmen Orozco-Mosqueda et al., 2013, Castulo-Rubio et al., 2015). *Arthrobacter agilis* UMCV2 significantly increased both *Medicago sativa* and *Sorghum bicolor* shoot biomass, as well as stimulated their iron scavenging systems (del Carmen Orozco-Mosqueda et al., 2013, Castulo-Rubio et al., 2015, Velázquez-Becerra et al., 2011). However, unlike BD237, UMCV2 did not increase root length or root weight (Fig. 2) (Velázquez-Becerra et al., 2011, Castulo-Rubio et al., 2015, del Carmen Orozco-Mosqueda et al., 2013).

The VOC mediated growth promotion of *B. distachyon* presented here is more subdued compared with previous studies, where a 3 – 6 fold increase in biomass has been reportedly induced by various microbial VOCs (Fig. 2a, c) (Blom et al., 2011a, Hao et al., 2016). These multi-fold increases in biomass have been reported in dicot plants such as *Arabidopsis* or *Nicotiana*. Studies using monocot plants report a similar, more subdued growth promotion effect as observed in the present study (Almeida et al., 2023, Delaplace et al., 2015).

*Bacillus* GB03 was the first bacterial strain reported to produce plant growth modulating VOCs, significantly increasing *A. thaliana* growth (Ryu et al., 2003). Since then, GB03 volatiles have been shown to stimulate as high as a 6-fold increase in other dicot plants such as *Nicotiana* (Zhang et al., 2007, Zhang et al., 2008a, Cappellari and Banchio, 2020). In monocots, *Bacillus* GB03 VOCs have a lesser growth promotion effect, suggesting that the same mVOCs have differential effects on dicots compared to monocots. In *B. distachyon,* GB03 VOCs elicited a comparatively lesser increase of 65% in shoot biomass, while no growth increase was observed in rice (Delaplace et al., 2015, Almeida et al., 2023). Thus, the current results are in agreement with previous studies where relatively mild growth promotion responses have been reported in monocots, compared to dicots. Additionally, it was found that soil grown *B. distachyon* plantlets exposed to BD237 mVOCs experience a 29% increase in dry shoot biomass (Fig. 3b). This result is not in agreement with a previous study which found that while mVOC mediated plant growth promotion can persist in soil, the beneficial effect is lessened (Cordovez et al., 2018). These results again highlight the divergent effect of mVOCs in dicots and monocots.

Previous works have highlighted the effect of bacterial growth medium on the plant growth modulation capacity of microbial VOCs (Blom et al., 2011a, Velázquez-Becerra et al., 2011, Fincheira and Quiroz, 2018, Fincheira et al., 2016). These studies found that in general, minimal media containing simple sugars stimulated the production of more plant growth promoting mVOCs than peptide rich media. Consequently, *B. distachyon* plantlets were exposed to mVOCs produced by *Pseudoarthrobacter* BD237 grown on either peptide rich LB agar medium or 1/2 MS media with glucose. Control plants grown with either LB or ½ MS media (no bacteria) were similar in their above and belowground biomass, root length and number of adventitious roots, indicating that bacterial growth media alone does not affect *B. distachyon* growth (Fig. 2). While LB media led to a significant increase in plant biomass, MS media led to a non-significant decrease in biomass (Fig. 2). These results contrast with a previous comprehensive study, which found that across 42 phylogenetically diverse soil bacteria, none produced plant growth inhibiting VOCs when grown on MS media (Blom et al., 2011a). However, as we only characterised the VOCs produced by BD237 on LB media and not MS media, it is unclear which mVOCs contributed to the observed plant growth reduction. Collectively, these results strengthen the notion that the plant growth modulating effect of mVOC are dependent on bacterial media (Ryu et al., 2020, Groenhagen et al., 2013).

VOCs produced by BD237 grown on LB media were identified via SPME-GC-MS (Table 4.1). Most detected VOCs have not been detected in previously reported *Pseudoarthrobacter* volatilomes, except for benzyl alcohol, phenylethyl alcohol and 3-methyl-butanol (del Carmen Orozco-Mosqueda et al., 2013, Velázquez-Becerra et al., 2011). Several of the detected VOCs have reported dicot plant growth modulating effects, while others have unknown effects on plant growth. For example, the current results are congruent with reports of 2 – nonanone improving *Oryza sativa* (rice)*, Lactuca sativa* (lettuce) and *Solanum lycopersicum* (tomato) growth, but are in contrast to those in *A. thaliana* growth, where growth was inhibited (Fig. 4) (Fincheira and Quiroz, 2019, Fincheira et al., 2020, Groenhagen et al., 2013, Almeida et al., 2023). The current results are also consistent with reports of benzyl and phenylethyl alcohols increasing *A. thaliana* growth (Fig. 4) (Groenhagen et al., 2013, Camarena-Pozos et al., 2019). Additionally, 2-heptanone has also been found to increase dicot growth, although a beneficial effect was not observed here (Fig. 4) (Jiang et al., 2019). Interestingly, the study by Camarena-Pozos and colleagues (2019) found that 3-methyl-butanol significantly increased *A. thaliana* and *Agave salmiana*, whereas this study found no growth stimulating effect of this VOC in *B. distachyon* (Fig. 4), suggesting that this mVOC may cause divergent responses in monocots and dicots.

### *Pseudoarthrobacter* BD237 mVOCs modulate the *B. distachyon* root transcriptome

Previously, VOC mediated growth promotion of dicots has been shown to result primarily from modulation of phytohormone networks, most notably through auxin, ethylene and cytokinin pathways (Zhang et al., 2007, Grossi et al., 2016, Hao et al., 2016, Gao et al., 2022, Ryu et al., 2003, Jiang et al., 2019). Thus, it was hypothesised that VOC mediated growth promotion in *B. distachyon* results from a similar modulation of its phytohormone pathways. To this end, an experiment was conducted where the transcriptomic response of *B. distachyon* roots after exposure to *Pseudoarthrobacter* BD237 VOCs was investigated.

*Pseudoarthrobacter*BD237 mVOCs caused a clear difference in gene expression between control and treatment samples (Fig.5) Subsequently, it was found that transcription of several key genes in the photosynthetic and carbon fixation pathways were significantly upregulated (Fig. 6, Supp. Table 2). Genes coding for RuBisCo (RBCS) and its smaller subunit RBCS1 were significantly upregulated in BD237 mVOC exposed roots compared to controls (Fig. 7c) These enzymes are crucial for photosynthesis and catalyse the primary reaction by which inorganic carbon is fixed in plants (Erb and Zarzycki, 2018). Phosphoribulokinase (PRK), the generator of substrate for RuBisCo was also significantly upregulated. PRK and RBCS are thought to be among the main determinants of a plant’s metabolic rate and growth due to their roles as key enzymes in carbon fixation (Miziorko, 2000). Transcription of chaperone enzymes such as RBCS2, required for proper assembly of RuBisCo, were also significantly upregulated (Saschenbrecker et al., 2007). The observed upregulation of key photosynthetic genes in the present work is congruent with past studies which observed a heightened photosynthetic rate in mVOC exposed plants (Fig. 7c) (Zhang et al., 2008b, Lin et al., 2025, Wang et al., 2023). However, certain photosynthesis related genes upregulated here, such as the chlorophyll binding protein LHCP (Fig.7c), were downregulated in an mVOC exposed dicot, suggesting that mVOC mediated modulation of photosynthesis many occur differently between monocots and dicots (Wang et al., 2023). Collectively, the present results suggest VOC exposed *B. distachyon* plantlets may have had a heightened photosynthetic rate leading to higher biomass, although this should be further experimentally confirmed. However, it should be noted that light exposure in the current experimental set up may have triggered root plastids exposed to mVOCs to be converted to chloroplasts and begin photosynthesising. It is possible that BD237 mVOCs also triggered photosynthesis in leaves, although this was not measured in the current study. In this way, observed enrichment of photosynthesis related genes may have been an artefact of the presently used methodology.

Several genes associated with a plant’s response to abscisic acid (ABA) and water deprivation were significantly upregulated (Fig. 6). ABA signalling plays an important role in a plant’s response to stressors. This was demonstrated with pea plants (*Pisum sativum*) defective in ABA biosynthesis and unable to close their stomata to preserve moisture in response to water deficit, as is usually observed in wild type plants (McAdam et al., 2015). Several key genes in the abscisic acid response pathway were significantly upregulated (Fig. 7b). For example, the catalase 1 (CAT1) gene was upregulated in VOC treated *B. distachyon* plantlets. In maize, CAT1 production is upregulated in response to abscisic acid and plays a crucial role in a plant’s response to stress by controlling cellular levels of hydrogen peroxide (H_2_O_2_) (Williamson and Scandalios, 1992). A 2021 study showed that mVOC exposed maize plants exhibited a higher drought tolerance, at least partially due to a significant increase in catalase content in treated plants (Yasmin et al., 2021). The dehydrin 3 (DHN3) gene was also upregulated in response to mVOC exposure and has also been shown to be upregulated in response to ABA application (Ding et al., 2020). Dehydrins are known to play important roles in plant response to drought and salinity stress, primarily via scavenging of oxygen radicals (Zhou et al., 2022). When overexpressed, these enzymes improve plant drought tolerance, while their silencing confers drought intolerance (Yu et al., 2019, Kumar et al., 2014). The MYB2 transcription factor was also significantly upregulated in mVOC exposed *B. distachyon* plantlets (Fig. 7b). The MYB family of transcription factors are known to play a role in a plants response to environmental stressors such as water deprivation (Urao et al., 1993). Transcription of MYB genes have previously been found to be modulated in response to mVOCs (Kim et al., 2015b, Hao et al., 2016, Shi et al., 2017, He et al., 2022, Paul et al., 2022). In one case, mVOC exposure also led to induced systemic resistance, suggesting a role of MYB factors in mVOC stimulated plant stress resistance (Zamioudis et al., 2015). A number of studies have found a similar protective effect against salt and drought stress elicited by mVOCs, albeit primarily in dicots and usually via modulation of salicylic or jasmonic acid pathways, rather than the abscisic acid pathway (Zhang et al., 2008a, Cho et al., 2008, Vaishnav et al., 2015, Ledger et al., 2016, Cappellari and Banchio, 2020, Liu et al., 2020). Similarly, studies focusing on monocots found that mVOC mediated abiotic stress tolerance was dependant on salicylic acid signalling (Mahadik and Kumudini, 2025). However, upregulation of stress response related genes may also have been a response to reactive oxygen species generated as photosynthesis by-products. For example, H_2_O_2_ is an important mediator of stress responses in cereals and is also produced during photosynthesis (Gill and Tuteja 2010). Taken together, the modulation of several key stress response genes in *B. distachyon* upon mVOC exposure suggests that BD237 volatiles may increase the resistance of *B. distachyon* to abiotic stressors via abscisic acid mediated signalling.

Conversely, genes linked to the nitrate assimilation pathway were significantly downregulated (Fig.7d, Table S3). These included high affinity nitrate transporters (NRT2.1, NRT2.3), a nitrite reductase (NIR1), a nitrate transporter activating protein (NAR2.1) and nitrate reductases (NIA1) (Fig.7d) with NRT2.1 and NAR2.1 observed as the most significantly downregulated genes (Fig. S4). Downregulation of NRT genes in response to mVOCs has been previously observed in tobacco, although this was not discussed by the authors (Kim et al., 2015b). Conversely, a study by Corodovez et. al (2018) found that mVOC exposure led to upregulation of NRT2.1 in *A. thaliana* roots, with the authors speculating that this gene may play a role in the regulation of lateral root growth (Cordovez et al., 2018). An additional study found that mVOCs lead to upregulation of NRT genes in *A. thaliana*, where the authors suggest that these genes may be required for communication between plants and PGPR (Lin et al., 2025).

The downregulation of nitrate metabolism genes observed in this study could be due to several reasons. Firstly, the stimulation of adventitious root growth in BD237 mVOC exposed plantlets likely led to an increased nutrient uptake. While not experimentally confirmed, the N status of mVOC exposed plantlets seemed sufficient, with no plantlets exhibiting visible signs of chlorosis (Fig. S2). NRT2.1 and NAR2.1 are required for activity of the inducible high-affinity nitrate transport system (IHATS) and are triggered when the plant is grown in low nutrient environments (Forde, 2000). Thus, if the plant has satisfied its N requirements, the IHATS and its associated genes may have been downregulated.

Secondly, nitrate acts not only as a vital plant nutrient, but also as a signalling molecule in the regulation of carbon metabolism (Scheible et al., 1997). As such, the modulation of genes related to nitrate metabolism here may have impacted photosynthesis. Subsequently, the downregulation of certain nitrate metabolism genes observed in this study may have contributed to the heightened expression of key photosynthetic genes rates, although this should be experimentally confirmed through gene knockout or silencing studies.

Finally, NRT2.1 has been shown to be a repressor of lateral root growth (Little et al., 2005). Thus, mVOC mediated downregulation of NRT2.1 observed here may have contributed to increased growth of adventitious roots. Additionally, as NRT2.1 requires the expression of NAR1 to effectively transport nitrate, their expression is tightly linked, where a reduction in expression of either would lead to a reduction in the other (Kotur et al., 2012).

### *B. distachyon* VOCs do not modulate the *Pseudoarthrobacter*BD237 root transcriptome

Exposure to *B. distachyon* VOCs did not signicantly alter the BD237 transcriptome, with no genes meeting the threshold of an adjusted p value <0.05 and |log₂FC| > 1). This result contrasts with a previous study where *Arabidopsis* emitted methyl jasmonate were shown to upregulate biofilm associated genes and downregulate motility associated genes in the model rhizosphere bacteria, *Pseudomonas protegens* Pf-5. The same study found that monocot and dicot root VOCs induced biofilm production in diverse rhizosphere microbial community, although the abundance of only around 9% of this community was altered upon VOC exposure, implying that many rhizosphere bacteria may not respond to plant VOCs, including *Pseudoarthrobacter* BD237. Instead, BD237 and similar microbes may constitutively emit bioactive VOCs.

## Conclusion

In conclusion, this study has isolated a *Pseudoarthrobacter* strain capable of increasing the dry biomass, root number and root length of the model grass *Brachypodium distachyon* via its VOCs. VOC production by BD237 was shown to be culture media dependent. The VOCs phenylethyl alcohol, benzyl alcohol and 2-nonanone identified via SPME-GC-MS were shown to similarly promote *B. distachyon* growth when applied as pure VOCs at different concentrations. Finally, while *B distachyon* VOCs did not elicit a transcriptomic response in BD237, root transcriptomic analysis showed that key genes in photosynthesis and stress response pathways were upregulated, while genes related to nitrate metabolism were downregulated upon exposure to BD237 mVOCs, with a possible link to stimulation of adventitious root growth. However, additional *in-planta* assays are required to confirm if mVOC can heighten photosynthetic rates and confer resistance to salinity and drought stress.

---

**Figure S1:**
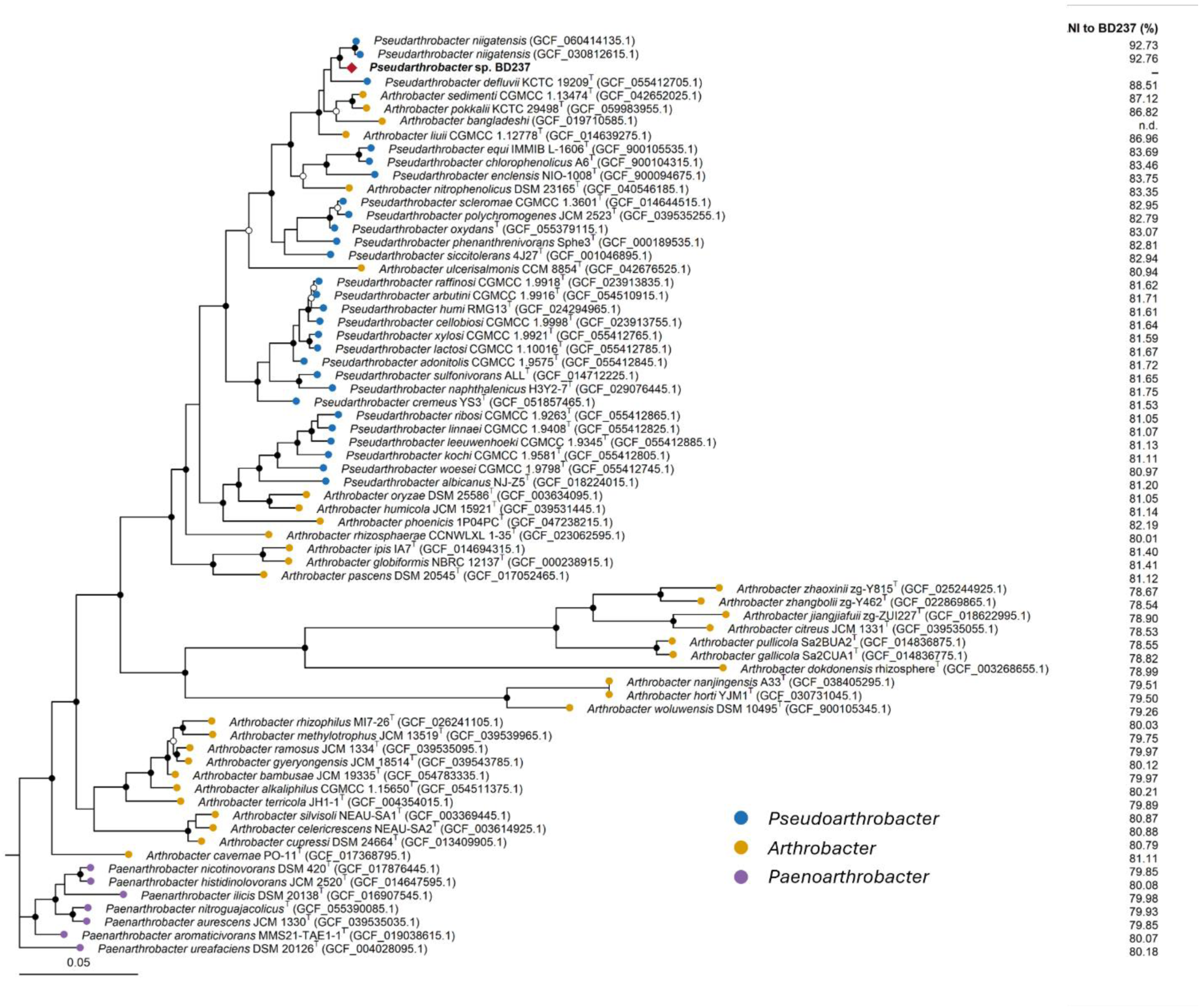
Maximum-likelihood phylogenomic tree of 69 genomes of *Pseudoarhtrobacter*, *Pseudoarthrobacter*and *Paenoarhtrobacter*, inferred in IQ-TREE from a concatenated alignment of 120 bacterial single-copy marker proteins (bac120) generated with GTDB-Tk. *Pseudoarthrobacter* sp. BD237 is shown in bold with a red diamond; remaining tip symbols are coloured by genus following LPSN, and a superscript T denotes a genome sequenced from type material. Node symbols indicate branch support: filled circles, SH-aLRT ≥ 80% and ultrafast bootstrap ≥ 95%; open circles, one of the two criteria met; unmarked nodes met neither criteria. The tree was rooted using *Paenarthrobacter* species. and the right column shows average nucleotide identity (ANI) to BD237 calculated with FastANI. The scale bar represents 0.05 amino-acid substitutions per site.

**Figure S2:**
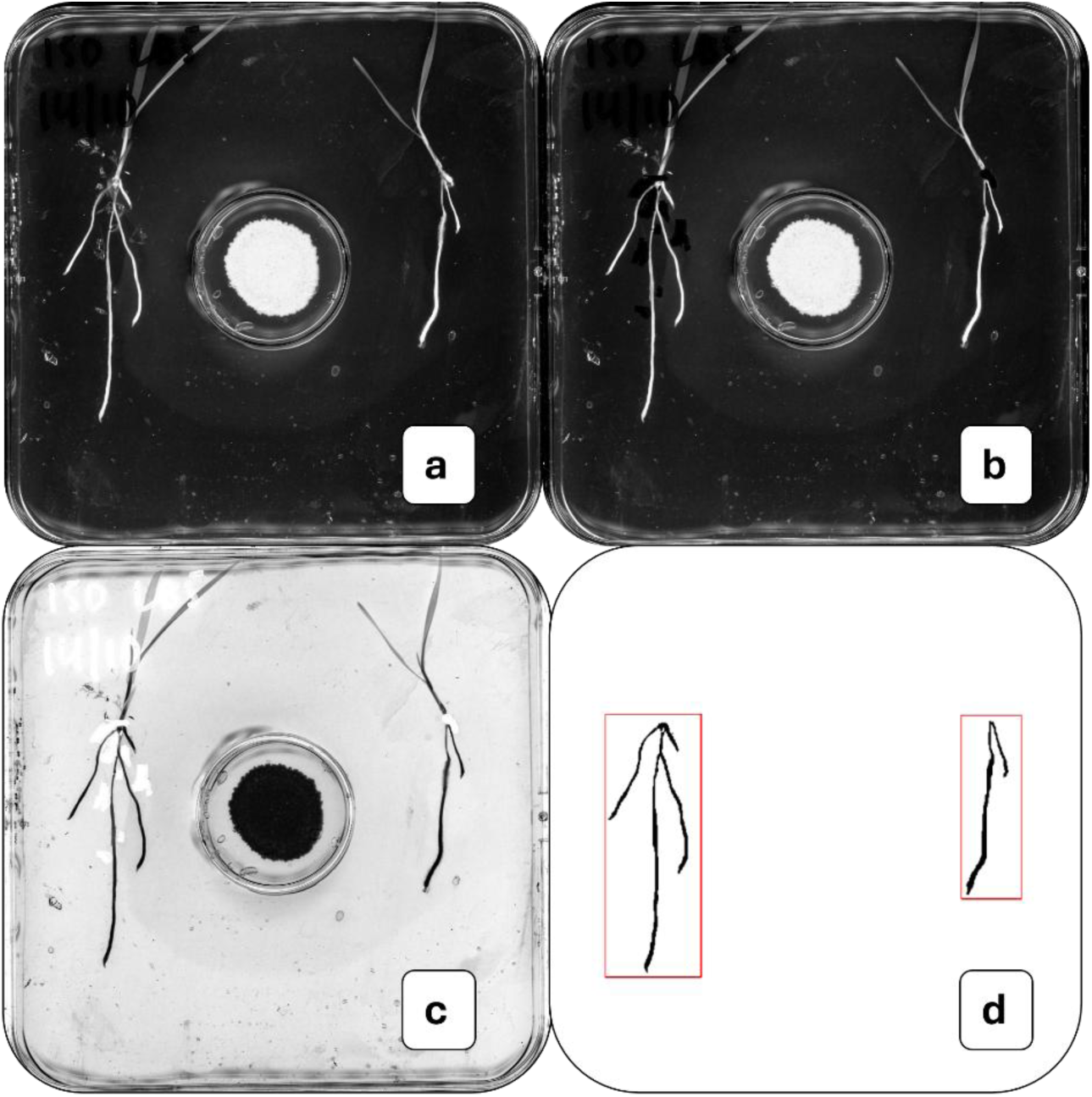
Processing and analysis of *B. distachyon* root scans. Raw scans (a) were manually edited with Microsoft Paint to remove larger image artefacts which interfere with downstream processing (b). Scans were then colour inverted (c) and roots sections manually specified (d) before root lengths were calculated.

**Figure S3:**
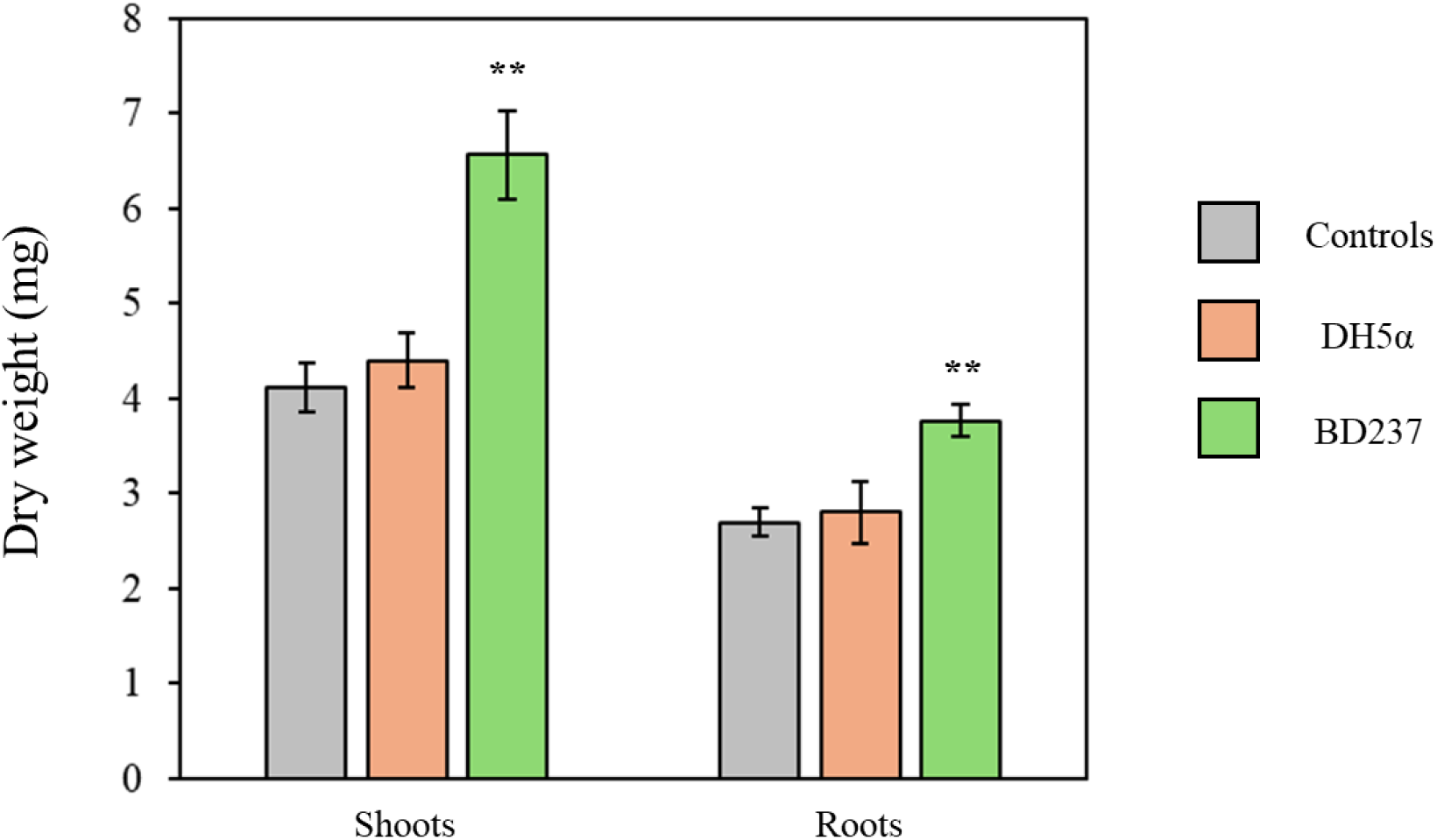
Shoot dry weight of *B. distachyon* plantlets exposed to *Pseudoarthrobacter*BD237 or *E. coli* DH5α mVOCs for 10 days. * = p < 0.05, ** = < 0.01. Significant differences were found via one-way ANOVA and post-hoc Tukey tests.

**Figure S4:**
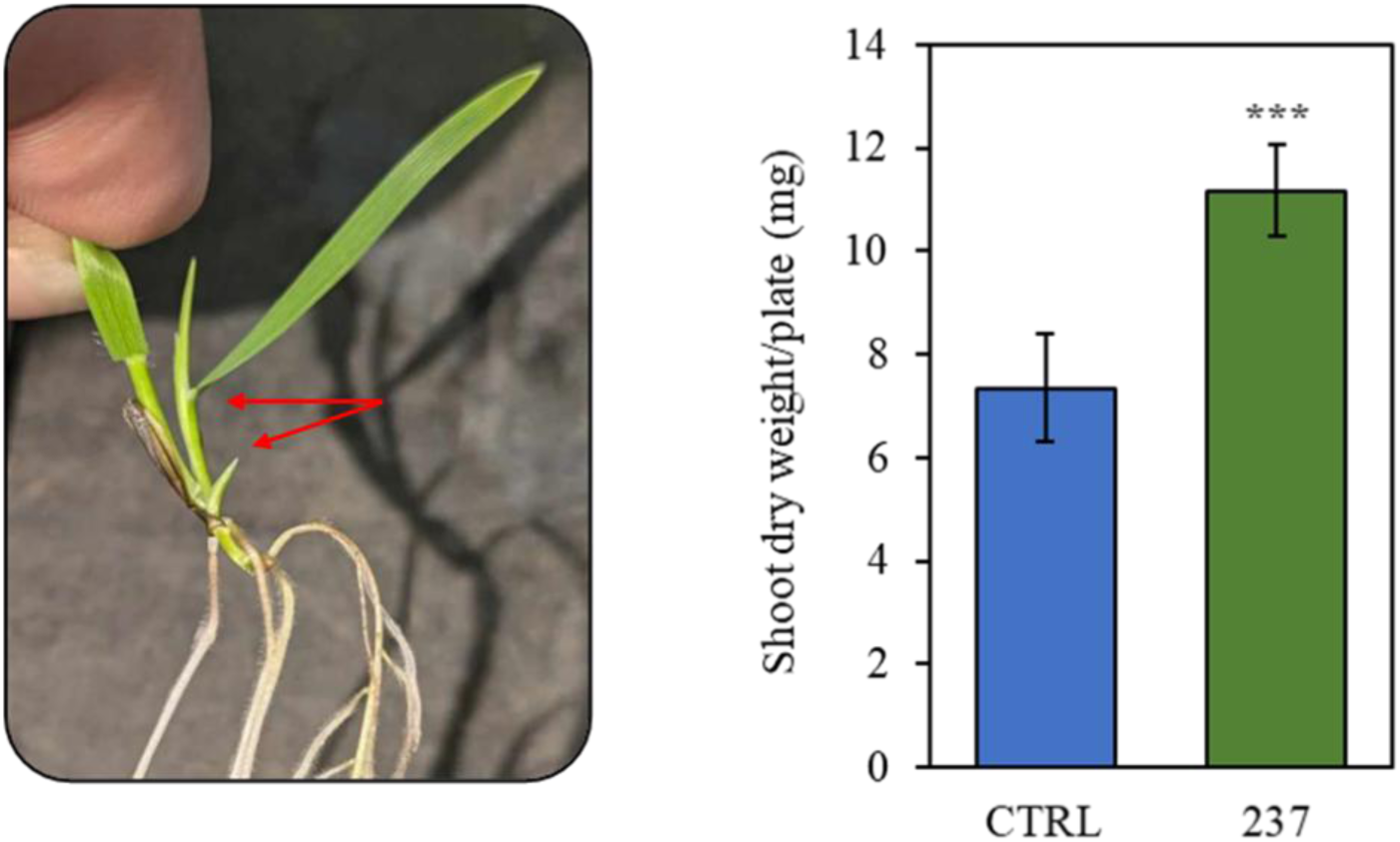
Dry shoot weight of *B. distachyon* plants exposed to *Pseudoarthrobacter* BD237 volatiles for 10 days (roots were harvested for transcriptomic analysis). Red arrows indicate tillering induced in *B. distachyon* by *Pseudoarthrobacter* BD237 volatiles. *** = p < 0.001. Significant differences were determined using unpaired T-tests.

**Figure S5:**
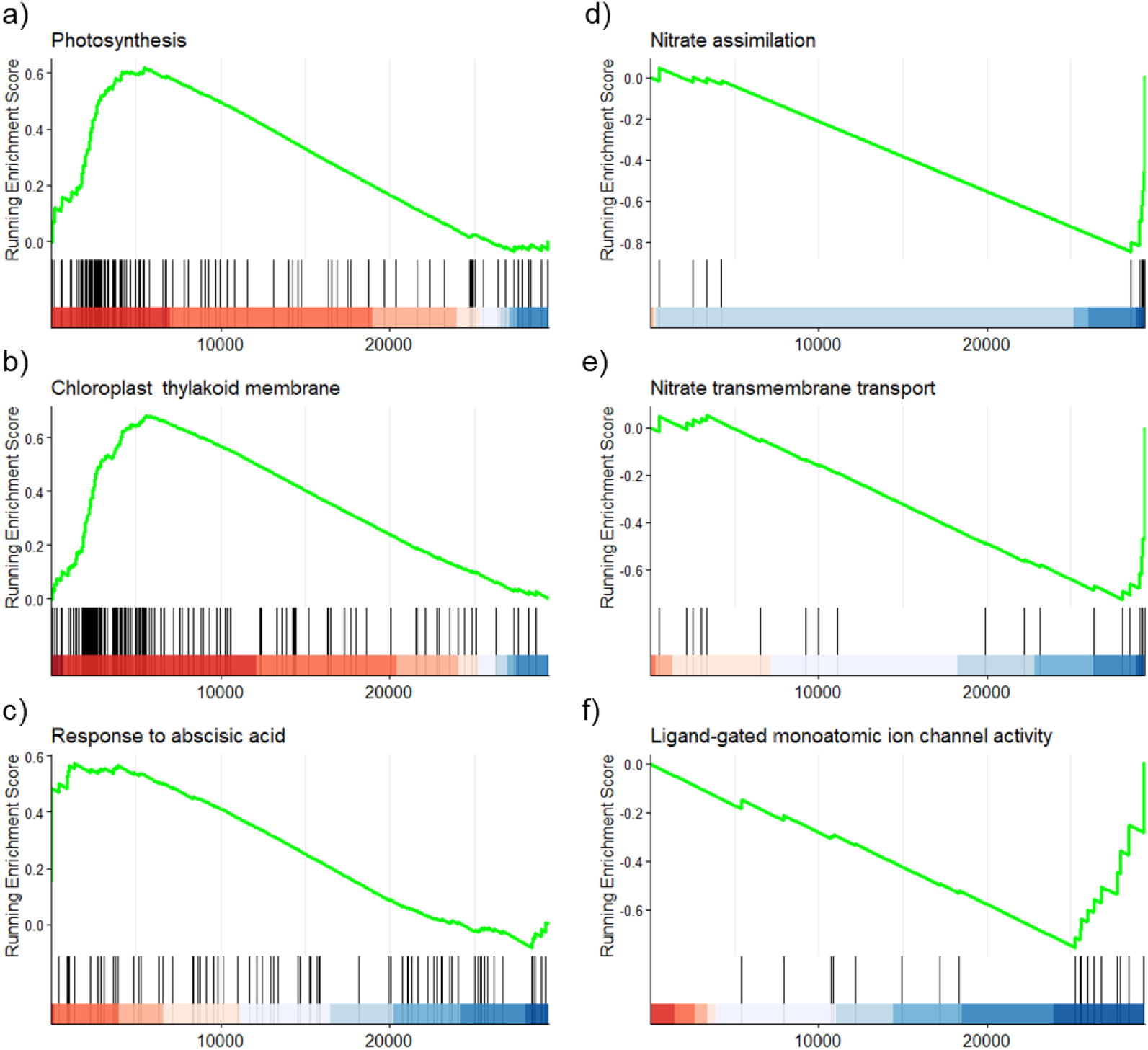
Normalised enrichment score (NES) plots of up (a, b & c) and down (d, e & f) regulated GO pathways, identified via GSEA analysis. These plots show where in the ranked list of genes (by fold change), individual genes related to pathways were placed.

**Figure S6:**
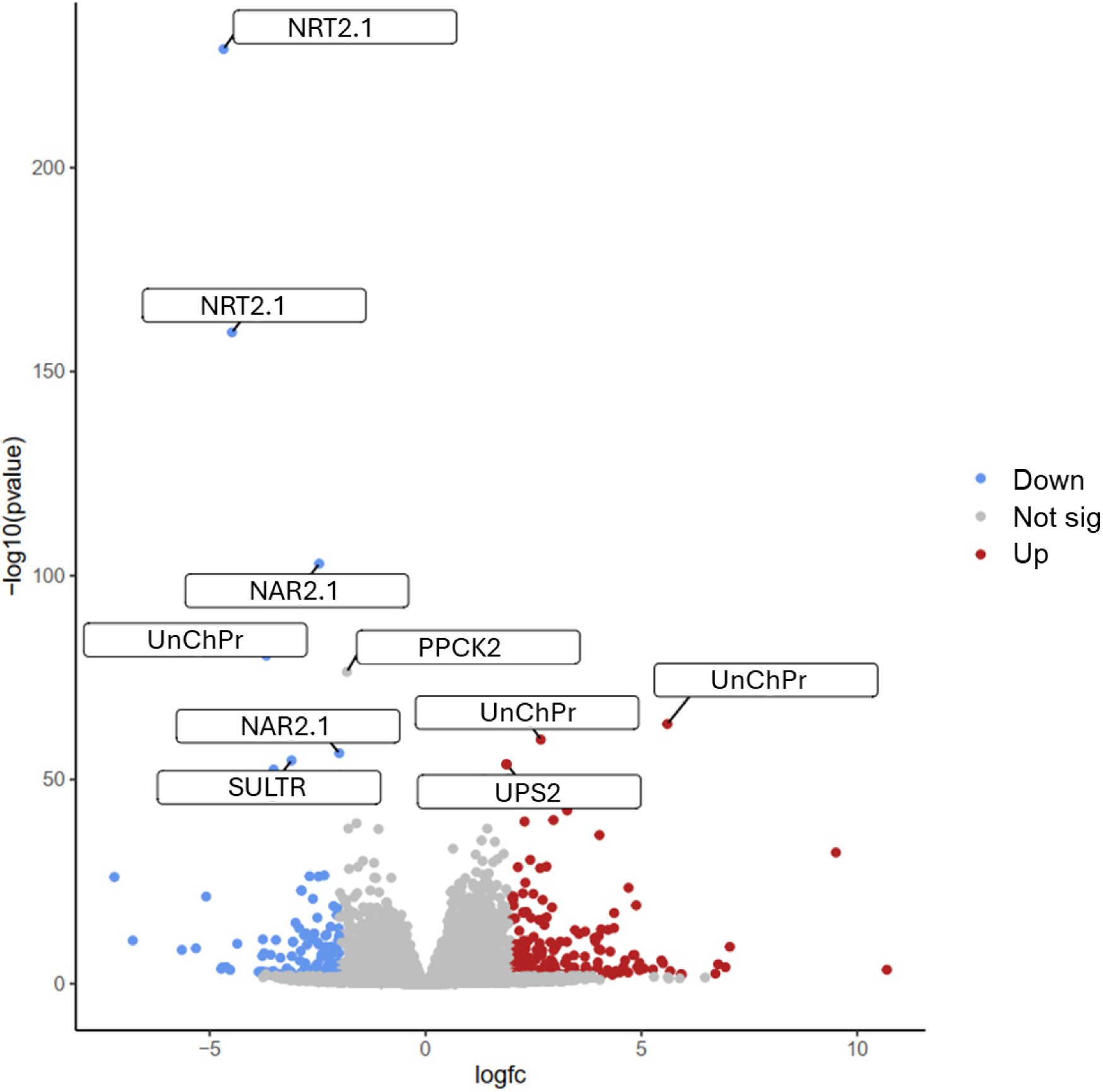
Volcano plot showing significantly up and down regulated genes in *B. distachyon* roots after exposure to BD237 mVOCs. Significantly differentially expressed genes were defined as those with a |log2(Fold Change)| > 1 (2-fold change) and padj < 0.05. NRT2.1 = High affinity nitrate transporter; NAR2.1 = High affinity nitrate transporter activating protein; UnChPr = Uncharacterised protein; SULTR = Possible sulfate transporter; PPCK2 = Phosphoenolpyruvate carboxylase kinase 2; UPS2 = Ureide permease 2.

**Table S1.** : *B. distachyon* RNAseq mapping and assignment rates

| Sample | HISAT2 mapping rate (%) | featureCounts assignment rate (%) |
| --- | --- | --- |
| PC1 | 97.2 | 82.7 |
| PC2 | 96.3 | 67.0 |
| PC3 | 97.0 | 79.3 |
| PC4 | 97.1 | 82.5 |
| PC5 | 96.5 | 83.9 |
| PT1 | 96.9 | 80.9 |
| PT2 | 96.6 | 76.9 |
| PT3 | 96.4 | 80.4 |
| PT4 | 96.9 | 78.7 |
| PT5 | 97.2 | 79.9 |
|  | AVG = 96.8% | AVG = 80.6% |
| Sample | HISAT2 mapping rate (%) | featureCounts assignment rate (%) |
| MC1 | 99.2 | 76.9 |
| MC2 | 99.4 | 79.1 |
| MC3 | 99.2 | 73.3 |
| MC4 | 99.3 | 78.4 |
| MC5 | 99.3 | 74.2 |
| MT1 | 98.8 | 76.4 |
| MT2 | 99.2 | 75.6 |
| MT3 | 98.9 | 75.4 |
| MT4 | 98.3 | 76.2 |
| MT5 | 99.1 | 68.3 |
|  | AVG = 99.1% | AVG = 75.4% |

Control samples = “PC/MC”, mVOC treated samples = “PT/MT”

**Table S2.**
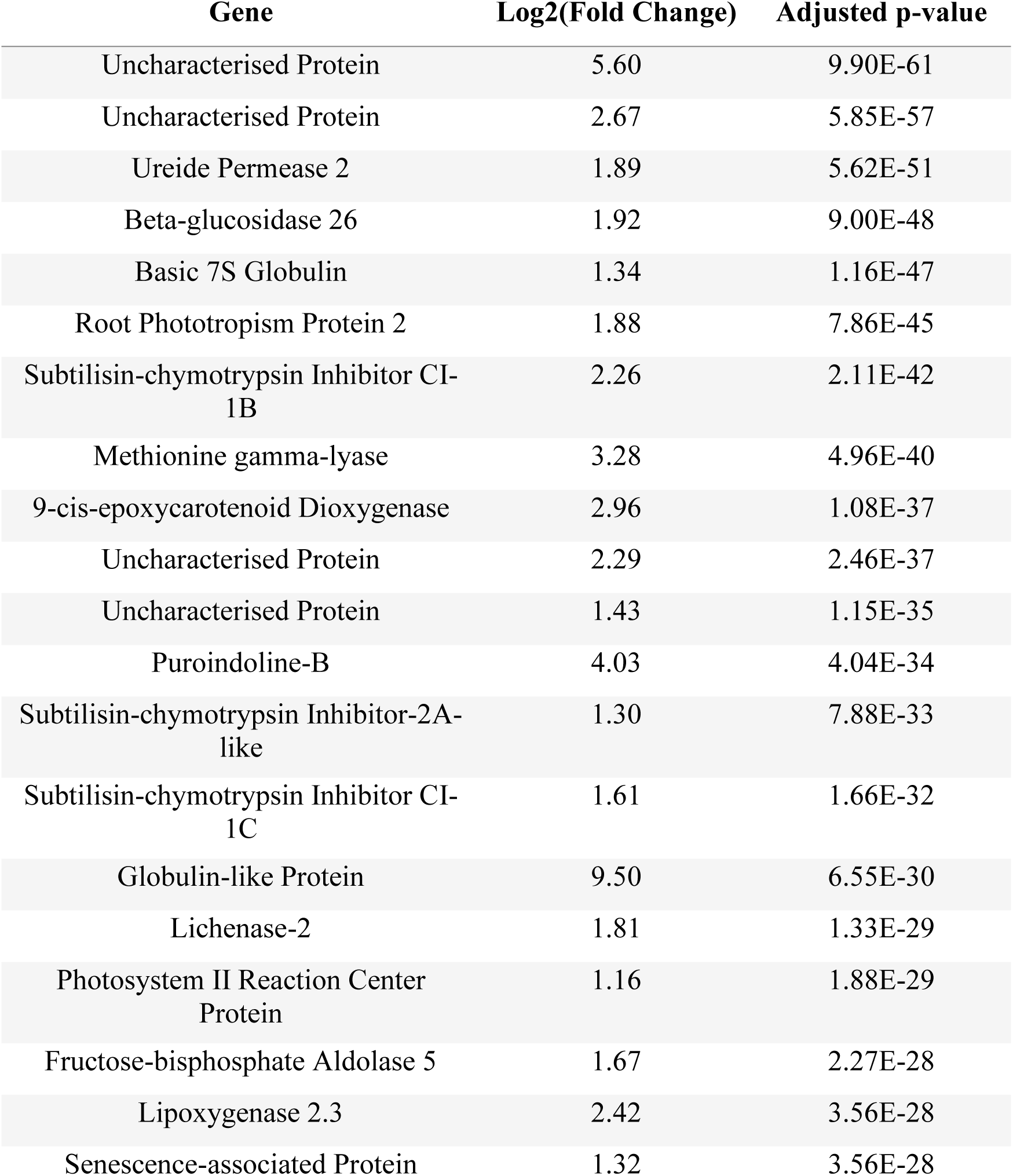
: Top 20 most significantly upregulated genes in *B. distachyon* roots upon exposure to *Pseudoarthrobacter* BD237 mVOCs

| Gene | Log2(Fold Change) | Adjusted p-value |
| --- | --- | --- |
| Uncharacterised Protein | 5.60 | 9.90E-61 |
| Uncharacterised Protein | 2.67 | 5.85E-57 |
| Ureide Permease 2 | 1.89 | 5.62E-51 |
| Beta-glucosidase 26 | 1.92 | 9.00E-48 |
| Basic 7S Globulin | 1.34 | 1.16E-47 |
| Root Phototropism Protein 2 | 1.88 | 7.86E-45 |
| Subtilisin-chymotrypsin Inhibitor CI-1B | 2.26 | 2.11E-42 |
| Methionine gamma-lyase | 3.28 | 4.96E-40 |
| 9-cis-epoxycarotenoid Dioxygenase | 2.96 | 1.08E-37 |
| Uncharacterised Protein | 2.29 | 2.46E-37 |
| Uncharacterised Protein | 1.43 | 1.15E-35 |
| Puroindoline-B | 4.03 | 4.04E-34 |
| Subtilisin-chymotrypsin Inhibitor-2A-like | 1.30 | 7.88E-33 |
| Subtilisin-chymotrypsin Inhibitor CI-1C | 1.61 | 1.66E-32 |
| Globulin-like Protein | 9.50 | 6.55E-30 |
| Lichenase-2 | 1.81 | 1.33E-29 |
| Photosystem II Reaction Center Protein | 1.16 | 1.88E-29 |
| Fructose-bisphosphate Aldolase 5 | 1.67 | 2.27E-28 |
| Lipoxygenase 2.3 | 2.42 | 3.56E-28 |
| Senescence-associated Protein | 1.32 | 3.56E-28 |

**Table S3.** : Top 20 most significantly downregulated genes in *B. distachyon* roots upon exposure to *Pseudoarthrobacter* BD237 mVOCs

| Gene | Log2(Fold Change) | Adjusted p-value |
| --- | --- | --- |
| High Affinity Nitrate Transporter 2.1 | -4.68 | 2.32E-225 |
| High Affinity Nitrate Transporter 2.1 | -4.48 | 3.01E-156 |
| High Affinity Nitrate Transporter-activating Protein 2.1 | -2.46 | 1.01E-99 |
| Uncharacterised Protein | -3.69 | 2.72E-77 |
| Phosphoenolpyruvate Carboxylase Kinase 2 | -1.82 | 1.98E-73 |
| High Affinity Nitrate Transporter-activating Protein 2.1 | -2.00 | 1.07E-53 |
| Sulfate Transporter 3.5 | -3.11 | 5.51E-52 |
| Uncharacterised Protein | -3.52 | 7.96E-50 |
| Indole-3-acetic acid-induced Protein | -1.98 | 2.30E-48 |
| Laccase 8 | -2.81 | 1.16E-47 |
| 2,3-bisphosphoglycerate-dependent Phosphoglycerate Mutase | -1.60 | 6.38E-37 |
| RING-H2 Finger Protein | -1.79 | 1.12E-35 |
| ACT Domain-containing Protein | -1.09 | 1.45E-35 |
| Auxin-induced Protein | -1.45 | 6.38E-28 |
| Tyrosine-sulfated Glycopeptide Receptor 1 | -1.19 | 1.77E-27 |
| Ubiquitin | -1.56 | 1.56E-26 |
| Zinc Finger Protein | -1.77 | 4.51E-26 |
| Basic Leucine Zipper | -2.34 | 1.62E-24 |
| Peroxidase 4 | -2.69 | 2.67E-24 |
| Uncharacterised Protein | -2.48 | 3.01E-24 |

**Table S4.** : Top 20 most significantly upregulated genes in *BD237* upon exposure to *B. distachyon* VOCs

| Gene | Log2(Fold Change) | Adjusted p-value |
| --- | --- | --- |
| TetR/AcrR Family Transcriptional Regulator | 0.57 | 1.69E-01 |
| GNAT Family N-Acetyltransferase | 0.44 | 1.97E-01 |
| RusA-like Resolvase | 1.01 | 2.22E-01 |
| Na<sup>+</sup>/H<sup>+</sup> Dicarboxylate Symporter (gltP) | 0.93 | 2.22E-01 |
| Mn-dependent Transcriptional Regulator MntR, DtxR Family (mntR) | 0.30 | 2.22E-01 |
| Nucleotidyltransferase Family Protein | 0.38 | 2.22E-01 |
| Fe(3<sup>+</sup>) Ions Import ATP-binding Protein FbpC (fbpC) | 1.92 | 2.22E-01 |
| Flagellar Biosynthesis Protein | 0.91 | 2.22E-01 |
| Holin | 1.25 | 2.22E-01 |
| 4-Carboxy-4-hydroxy-2-oxoadipate Aldolase/Oxaloacetate Decarboxylase (ligK) | 0.73 | 2.22E-01 |
| Na<sup>+</sup>/Melibiose Symporter-like Transporter | 0.73 | 2.22E-01 |
| Iron ABC Transporter Permease | 1.28 | 2.22E-01 |
| DNA-binding Transcriptional Regulator, GntR Family (mngR) | 0.25 | 2.22E-01 |
| Transcriptional Regulatory Protein | 0.30 | 2.22E-01 |
| MFS Transporter | 0.23 | 2.22E-01 |
| Iron ABC Transporter Substrate-binding Protein | 1.74 | 2.22E-01 |
| Oxidoreductase | 0.49 | 2.22E-01 |
| DUF6318 Domain-containing Protein | 0.79 | 2.22E-01 |
| DUF3072 Domain-containing Protein | 0.41 | 2.22E-01 |
| Secreted Protein | 2.48 | 2.22E-01 |

**Table S5.**
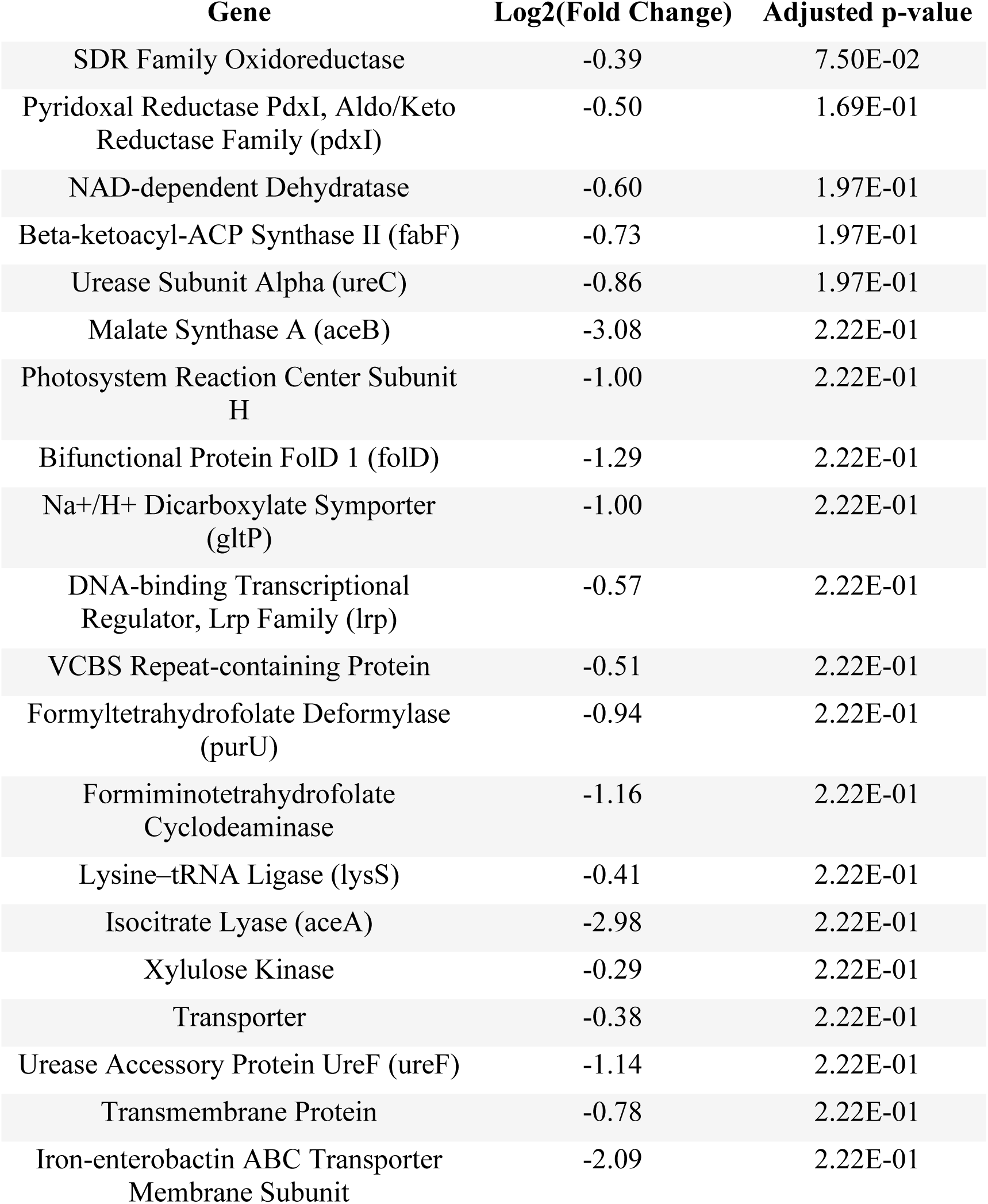
: Top 20 most significantly downregulated genes in *BD237* upon exposure to *B. distachyon* VOCs

